# Predictive all-atom simulations of disordered proteins and biomolecular condensates through osmometry-guided force-field optimization

**DOI:** 10.64898/2026.08.25.747127

**Authors:** Miloš T. Ivanović, Valentin von Roten, Benjamin Schuler, Robert B. Best

## Abstract

All-atom simulations with explicit solvent can provide a detailed and accurate description of dynamics and mechanisms in biomolecular systems, including intrinsically disordered proteins (IDPs) and their condensates. However, interactions involving charged residues and ions remain a persistent source of systematic error. Here we introduce an osmometry-guided optimization strategy that directly targets residue–residue, residue–ion and ion–ion interactions. Osmotic pressure provides key experimental information on molecular interactions and can be calculated directly and rapidly from simulations, enabling iterative force-field optimization. The resulting parameters improve agreement with single-molecule FRET data for IDPs, NMR relaxation data for an IDP–folded-domain complex, and chain dynamics and dimensions in biomolecular condensates of charged IDPs. For such condensates, simulations with our osmometry-optimized force field provide the missing link for predicting condensate dynamics across length and time scales. The strategy is broadly extensible to other interaction classes, including those governing protein–nucleic-acid assemblies.

---

Intrinsically disordered proteins (IDPs) and intrinsically disordered regions (IDRs) are central to signaling, transcription, molecular recognition and the organization of cellular assemblies.^1^ Rather than adopting one stable structure, they populate heterogeneous conformational ensembles whose populations and dynamics respond to sequence, binding partners, post-translational modification and solution conditions. These properties allow an IDR to recognize several partners, couple binding to regulation and remain dynamic within high-affinity complexes.^1–3^ IDPs and IDRs are also major constituents of biomolecular condensates, concentrated and dynamic assemblies formed via multivalent interactions between proteins and nucleic acids,^4^ which have been implicated in signaling, ribosome biogenesis, genome organization, transcription and stress response.^1,5,6^ The broad structural ensembles populated by IDPs make them difficult to model, and agreement with individual average observables does not ensure that the underlying conformational ensemble, contacts or timescales are correct.^7^ No single experimental technique fully resolves such conformational landscapes at atomistic resolution, making accurate molecular models essential for interpreting and predicting these ensembles.^8^ In condensates, this challenge is amplified because these properties depend not only on intramolecular interactions but also on many transient intermolecular contacts in a concentrated, often multicomponent environment.^9^

Coarse-grained models have long been the workhorse of condensate simulations because they can reach the system sizes and statistical sampling needed to calculate phase diagrams, compare sequences and investigate material-property trends.^10–13^ However, owing to the many degrees of freedom omitted, the time scales of dynamics in coarse-grained simulations are generally not accurate and can only approximately be rescaled for comparison with experiment.^9,12,14^ All-atom explicit-solvent force fields are able to play a complementary role to coarse-grained models: once optimized, the same model can be applied without system-specific parameter adjustment to monomeric IDPs, protein complexes and condensates, including systems containing both disordered and folded proteins, and the resulting dynamics can be directly compared with experiment on an absolute time axis.^12,15–17^

Recent advances in computer hardware and molecular-dynamics software are making all-atom condensate simulations increasingly feasible. For example, systems containing several million atoms can now be followed for several microseconds,^12,15–17^ but it is still critical to have an accurate energy function, or force field: because of their disorder, IDRs can substantially shift their configurational ensembles in response to even small force field errors,^18^ in contrast to the more robust situation for folded proteins.^19,20^ Over the past decade, protein force fields have improved markedly through refinements of backbone and side-chain torsions,^19,21,22^ protein–water interactions^20,23–26^ and, in some models, electronic polarization.^22,27^ These developments have enabled equilibrium folding calculations^28^ and realistic simulations of both folded and disordered proteins.^25,29,30^ However, interactions between charged groups have remained a persistent weakness of the widely used pairwise-additive models.^31,32^ Association measurements on side-chain analogues and NMR estimates of salt-bridge populations indicate that oppositely charged groups are too strongly associated in simulation.^31,33^ In a sequence-diverse set of 16 IDRs, deviations between simulated and experimental mean Förster resonance energy transfer (FRET) efficiencies were most strongly correlated with the fraction of charged residues and the number of possible salt bridges.^34^

Including explicit electronic polarizability would provide a more physically realistic form for Coulomb interactions, but polarizable force fields remain four-to ten-fold more expensive and are not yet routine for the multi-million-atom, multi-microsecond calculations considered here.^35^ Two practical corrections to interactions between charged groups have been successfully used within pairwise-additive models: Electronic-continuum-correction approaches scale ionic or charged-group partial charges to represent electronic screening implicitly.^36–39^ Alternatively, pair-specific Lennard-Jones terms instead rebalance selected short-range interactions without changing the integer net charges of residues or ions.^40^

Identifying accurate corrections requires experimental benchmarks that can identify the interaction responsible for a force-field deficiency. Rather than attempting to optimize a force field directly in a protein or condensate, where many interaction types are inseparably coupled, we isolate each charge interaction class in a minimal solution that retains the relevant chemical groups. Osmotic pressure is an observable particularly suited to this strategy because it can be measured for defined solutions and calculated directly and rapidly for the same solution composition in simulation.^40–42^ Here we thus develop an osmometry-guided workflow to optimize charge interactions in an all-atom force field (Fig. 1). We augment the limited available osmotic pressure data for monovalent salts and amino acids with additional, tailored experiments targeting residue–residue, residue–ion and ion–ion interactions. We derive two force-field parameter sets that each reproduce the experimental observations: one based primarily on charge scaling and another that preserves integral charges while modifying Lennard-Jones interactions. We assess the optimized parameters against a large set of experimental data: single-molecule FRET measurements for 16 sequence-diverse IDRs; NMR relaxation data for a highly charged complex between an IDP and a protein containing a folded domain; and single-molecule FRET and nanosecond fluorescence correlation spectroscopy (nsFCS) measurements for four multicomponent condensates formed by charged disordered proteins. Residue-specific Lennard-Jones optimization yields a consistent overall improvement against this validation data set. Together, these results demonstrate that forcefield parameters optimized using osmotic pressures of simple amino acid and ion solutions transfer without further adjustment to proteins and multicomponent condensates. Because the solutes can be selected to target the interaction class of interest, the approach provides a simple and broadly applicable experiment-based strategy for optimizing problematic force field interactions. This enables quantitative all-atom predictions of structure and dynamics across complex biomolecular systems.

**Figure 1:**
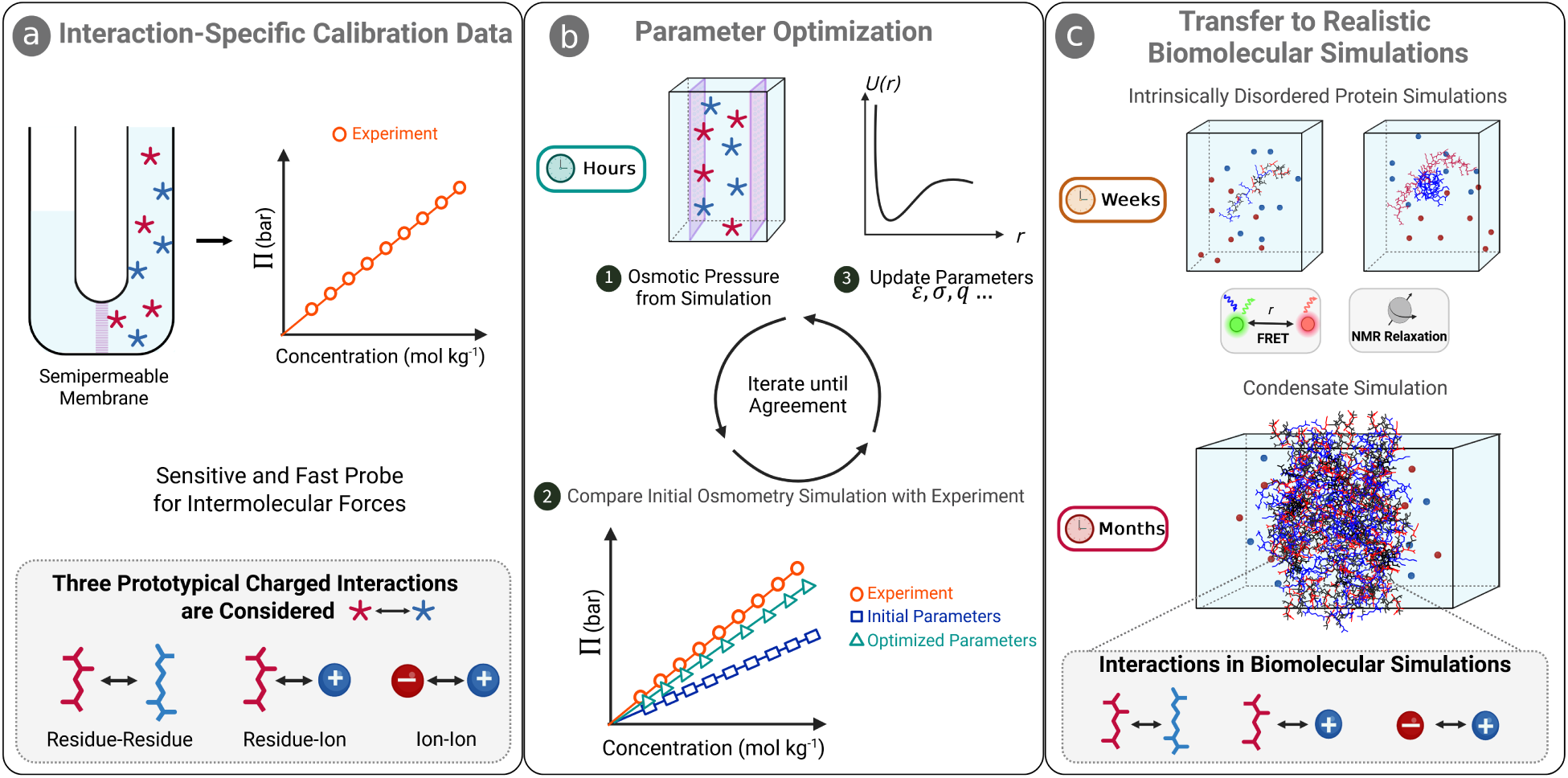
Osmometry-guided optimization and validation of charge interactions in all-atom force fields. **a,** Osmotic pressure, Π, is measured for defined solutions that isolate residue–residue, residue– ion and ion–ion interactions. The schematic illustrates the thermodynamic origin of osmotic pressure and does not depict the experimental measurement setup. Blue and red denote positively and negatively charged species, respectively. Star-shaped symbols represent generic charged solutes, water is light blue. **b,** Comparison between simulated and experimental concentration-dependent osmotic pressures identifies interactions that are too attractive or too weak. In simulations, osmotic pressure can be calculated from the total force exerted on virtual semipermeable walls that mimic an osmotic membrane. Nonbonded parameters controlling Lennard-Jones interactions (*σ* and *ɛ*), and partial charges of atoms (*q*) are varied iteratively until the benchmark is reproduced. **c,** Final parameters are transferred without further adjustment to explicit-solvent simulations of IDPs, protein complexes and condensates and validated against FRET, NMR and nsFCS. The osmometry simulations converge within hours, whereas the production simulations of large systems require weeks to months. Molecular sizes are schematic and not drawn to scale.

## Results

### Osmotic pressure as a sensitive measure of molecular interactions

We split force-field optimization into calibration and validation stages that differ substantially in molecular and computational complexity (Fig. 1). For calibration, osmotic pressures are measured for solutions chosen to isolate classes of interactions between charged moieties that recur in biomolecular simulations (Fig. 1a). The present set comprises oppositely charged amino acid pairs, charged amino acids with monovalent counterions, and NaCl or KCl. These mixtures retain the relevant chemical groups while avoiding the conformational complexity of a protein and the coupled interaction network of a condensate. Nonetheless, the side chains already occur in the context of an amino acid, avoiding the need to assume transferability from small-molecule analogues.^43^

The experimental compositions are reproduced in explicit-solvent simulations with semipermeable walls (Fig. 2a–c), and the osmotic pressure is calculated from the mean restoring force acting on the confined solutes.^40^ Experiment and simulation can therefore be compared directly through the concentration dependence of the same thermodynamic observable. Because these calibration systems are small, candidate electrostatic and Lennard-Jones parameters can be varied iteratively until the simulated osmotic pressures reproduce the experimental measurements (Fig. 1b). Finally, the optimized parameters are applied without further modification to increasingly complex biomolecular systems and validated against multiple experimental observables (Fig. 1c).

**Figure 2:**
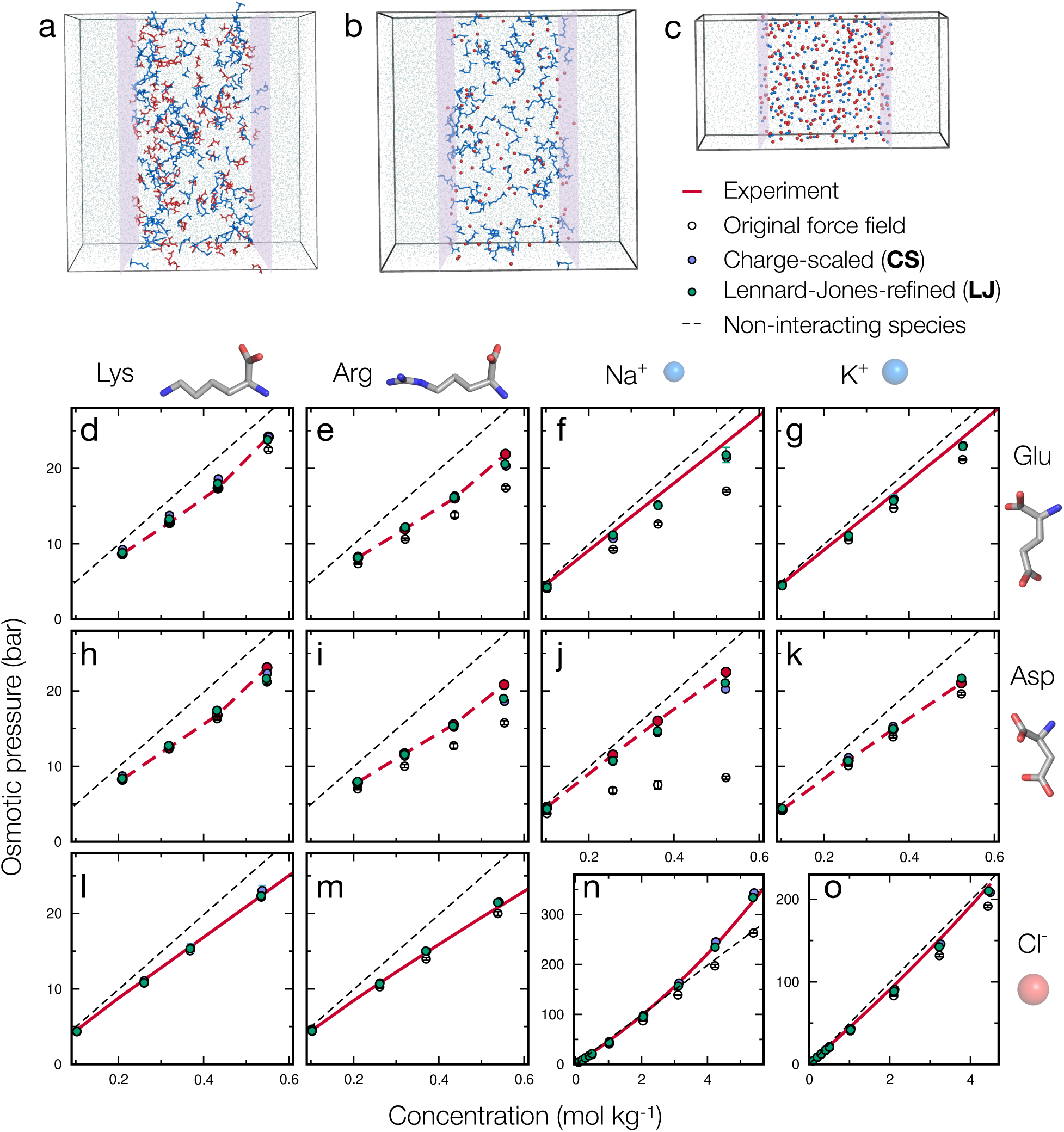
Osmometry benchmark and optimization of charge interactions. **a–c,** Representative simulation systems isolating residue–residue, residue–ion and ion–ion interactions. Positive species are blue, negative species red, water oxygens light blue and virtual semipermeable walls purple; hydrogen atoms are omitted. **d–o,** Interpolated experimental osmotic pressures (red lines) are compared with corresponding values calculated using three force fields: the original Amber ff99SBws (black circles), charge-scaled Amber ff99SBws-CS (purple circles), and Lennard-Jones-refined Amber ff99SBws-LJ (green circles). Dashed red lines denote vapor-pressure-osmometry measurements obtained in this work for Lys–Glu, Lys–Asp, Arg–Glu, Arg–Asp, Asp–Na^+^ and Asp–K^+^; solid red lines denote values taken from the literature. ^44–46^ Simulated osmotic pressures below the corresponding experimental values indicate excessive net attraction. The black dashed lines indicate the dependence expected for non-interacting species. Experimental and simulation uncertainties are defined in Methods. Most error bars are smaller than the plotting symbols.

#### A benchmark dataset resolves the relative strengths of charge interactions

For a solution of non-interacting solute species, the ideal osmotic pressure is Π_ideal_ = *c*_tot_*RT*, where *c*_tot_ = Σ*_i_ c_i_* is the total molar concentration of all solute particles, with each dissociated ionic species counted separately, *R* is the molar gas constant, and *T* is the absolute temperature. Attractions between solute molecules tend to lower Π relative to Π_ideal_, whereas repulsive interactions have the opposite effect. The osmotic coefficient, *ϕ* = Π*/*Π_ideal_, thus reports the net balance of solute–solute and solute–solvent interactions. We measured concentration-dependent osmotic pressures for equimolar Lys–Glu, Lys–Asp, Arg–Glu and Arg–Asp mixtures, and for Asp–Na^+^ and Asp–K^+^ solutions. Previously published values for the remaining residue–ion and ion–ion pairs complete our benchmark dataset.^44–46^ Together, the data cover the basic and acidic side chains and the monovalent ions used throughout the subsequent protein simulations. Measurements over a range of concentrations are important because a simulation may agree with experiment near the dilute limit yet deviate at high concentrations where repulsive interactions accumulate.

The measurements reveal substantial chemical specificity (Fig. 2). KCl and KGlu remain close to ideal over much of the investigated range, whereas most other pairs have osmotic coefficients below one, consistent with net attraction. At high concentration, NaCl instead reaches *ϕ >* 1. The strength inferred for amino acid association follows Arg–Asp *>* Arg–Glu *>* Lys–Asp *>* Lys–Glu (Fig. S1a). For amino acid–ion pairs, the corresponding order is Arg–Cl*^−^ >* Asp–K^+^ *>* Lys–Cl*^−^ >* Asp–Na^+^ *>* Glu–Na^+^ *>* Glu–K^+^ (Fig. S1b). The pronounced Arg vs. Lys difference is consistent with stronger Arg-dependent interactions reported in IDPs and condensates^16,47,48^ and with charge-patterned IDPs, where replacing Lys by Arg strongly affects salt-dependent chain expansion whereas replacing Glu by Asp has a much smaller effect.^49^ These chemically resolved measurements therefore provide a benchmark for testing whether a force field reproduces both the relative and overall strengths of charged interactions.

We calculated the corresponding osmotic-pressure curves based on simulations with Amber ff99SBws and TIP4P/2005s water, using the Luo–Roux monovalent-ion parameters as the starting point.^20,40,50^ Because standard Amber protein force fields do not provide a parameter set for isolated amino acids with both termini charged, we derived a shared zwitterionic backbone while retaining the original side-chain charges (Methods). Simulations of zwitterionic alanine and glycine agreed with literature data^51,52^ (Fig. S2) without adjustment, validating the terminal-group model. The original force field reproduced several interaction pairs reasonably but underestimated Π for several combinations, indicating excessive net attraction (Fig. 2). The largest deviations involved arginine and sodium with anionic amino acids, and Asp–Na^+^ showed the strongest discrepancy. This ordering is consistent with earlier measurements and calculations identifying overly favorable salt bridges in additive force fields.^31,33^ It also mirrors the stronger reweighting required for arginine-containing salt bridges inferred in our previous study.^34^

### Two optimization routes reproduce the osmotic-pressure benchmark

We explored two force-field optimization routes that alter the balance of solute–solute and solute–water interactions by prioritizing changes to different terms in the potential energy function. The first route weakens Coulomb interactions by downscaling the charges of charged residues and ions, followed by limited ion–water Lennard-Jones adjustments. The second route preserves the integral net charges of residues and ions and more extensively modifies Lennard-Jones interactions.

#### Route 1: Charge scaling correction

Following the rationale of electronic-continuum-correction models,^36–39^ we scaled the partial charges on the heavy atoms of charged side chains so that each side chain carried a reduced net charge of ±*q* rather than ±1*e*. We scanned *q* using the four amino acid-pair datasets at four concentrations and evaluated the global discrepancy from experiment. A charge magnitude of 0.93*e* minimized the unreduced *χ*^2^ (Fig. S3), reaching a value of 7.1 from an initial value of 45.6. We therefore scaled net charges of charged residues by 0.93 and applied the same scaling to all monovalent ions to preserve charge neutrality; this value is in a similar range as obtained in several earlier studies.^26,38,53,54^

In addition to the residue–residue data, charge scaling also brought the residue–ion and ion–ion osmotic pressures closer to experiment, but interactions of Na^+^ and K^+^ with acidic residues remained too attractive, whereas Cl*^−^* association with basic residues became slightly too weak. A single electrostatic scaling factor does not permit further independent tuning of pair interactions. To further improve residue–ion and ion–ion parameters, we therefore adjusted the Lennard-Jones diameter *σ* between inorganic ions and the water oxygen atom (OW), as related corrections have previously been required in charge-scaled models.^55,56^ Relative to values from the standard combination rule, the selected scaling factors were 0.965 for *σ*_Na+–OW_, 0.98 for *σ*_K+–OW_ and 1.005 for *σ*_Cl_*_−_*_–OW_ (Fig. S4). The small *σ*_Cl_*_−_*_–OW_ change had little effect on the already reasonable Lys–Cl*^−^* and Arg–Cl*^−^* curves. As an additional experimental constraint on ion pairing, we used Na^+^–Cl*^−^*and K^+^–Cl*^−^* coordination numbers determined by solution X-ray diffraction at near-saturating salt concentrations.^57^ These values quantify the average number of oppositely charged ions in direct contact, as opposed to ions separated by at least one hydration shell, and are 0.3 ± 0.1 for NaCl and 0.6 ± 0.1 for KCl.^57^ Together with the cation–water adjustments, this enabled the NaCl and KCl osmotic pressures and coordination numbers to be described with the same parameter set. The set of parameters optimized via this route was denoted Amber ff99SBws-CS.

#### Route 2: Lennard-Jones optimization with integral charges

The second route retained the formal net charges of all residues and ions. Instead, we first changed *σ* for interactions between OW and the heavy atoms of charged amino acid side chains. A uniform 1% reduction across the four charged residues lowered the global *χ*^2^ for the four residue-pair datasets from 45.6 to 6.0 (Fig. S5). Residue-specific optimization reduced it further, to 4.5. The selected *σ*_residue–OW_ scaling factors were 0.990 for lysine, 0.985 for arginine, 0.990 for aspartate and 0.995 for glutamate. Modifying *σ*_residue–OW_ shifts the balance between hydration and direct residue association while leaving formal charge and long-range electrostatics unchanged.

These modest changes in *σ*_residue–OW_ were sufficient for Lys–Cl*^−^* and Arg–Cl*^−^*, and improved, but did not fully correct, the interactions of acidic residues with Na^+^ or K^+^ (Fig. S7). We therefore also modified the ion Lennard-Jones parameters. Across commonly used monovalent-ion parameter sets, *σ* and the well depth *ɛ* differ surprisingly widely but are strongly anticorrelated (Fig. S6a), likely because of the emphasis on parameterization using solvation free energy, which is strongly dependent on the ion radius.^58^ Defining the effective radius *r*_eff_ as the distance at which the ion–water Lennard-Jones energy reaches 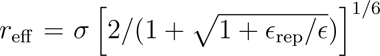, we identified a *σ*–*ɛ* relationship across several standard force-field parameter sets (Fig. S6d). Guided by this relationship, we varied *σ* and *ɛ* while changing *r*_eff_ by less than 2%. Increasing *σ*_K+_ weakened excessive Glu/Asp–K^+^ association, whereas lowering *ɛ*_K+_ partly opposed this change by shifting the competition between ion–water and residue–ion interactions. A 1.3-fold increase in *σ*_K+_ together with a 0.02-fold scaling of *ɛ*_K+_ best reproduced both acidic-residue datasets (Fig. S7c,d). For Na^+^, the residue–water changes already reduced excessive Glu/Asp–Na^+^ attraction. A further 1.3-fold increase in *σ*_Na+_ together with a 0.025-fold scaling of *ɛ*_Na+_ improved the agreement, and setting *σ*_Na+–OW_ to its original force-field value gave the best result within this functional form (Fig. S7e,f). Asp–Na^+^ remained somewhat too attractive, but further changes produced little improvement.

We finally tested ion–ion association, because changes made to improve residue–ion interactions can disrupt electrolyte behavior. Indeed, the modified K^+^ parameters overstabilized KCl at high concentration and increased the K^+^–Cl*^−^* coordination number to 2.92 ± 0.03, far above the experimental value of 0.6 ± 0.1.^57^ Pair-specific K^+^–Cl*^−^* parameters, with *σ*_K+–Cl*−*_ and *ɛ*_K+–Cl_*_−_* scaled by factors of 1.1 and 0.29, respectively, reproduced the KCl osmotic-pressure curve and yielded a coordination number of 0.50 ± 0.03, within error of the experimental value. The modified Na^+^ parameters had the opposite effect and made NaCl association too weak. Scaling *ɛ*_Na+–Cl_*_−_* by 0.29 restored agreement with the osmotic-pressure data and increased the Na^+^–Cl*^−^* coordination number from 0.0006 to 0.22±0.03, close to the experimental value of 0.3±0.1.^57^ These tests ensure that the improvement in amino-acid–ion interactions remains consistent with the experimental ion pair distribution. The complete set of parameters optimized via this route was denoted Amber ff99SBws-LJ.

### Parameter testing in folded and disordered proteins

Altogether, charge scaling (CS) and Lennard-Jones (LJ) optimization both reproduce the calibration data set, but what are the consequences of these parameter changes for the transferability to large biomolecular systems? The parameters optimized in the osmometry approach were transferred without further adjustment to biomolecular systems at different levels of complexity: monomeric IDRs, a folded–disordered protein complex and multicomponent condensates.

#### Improved consistency with FRET data for 16 disordered proteins

To determine whether agreement in the calibration systems transfers to proteins, we simulated 16 naturally occurring IDRs of identical length but diverse composition and charge patterning.^34,59^ Their mean transfer efficiencies, which report on the average dye-to-dye distances, had previously been measured by single-molecule FRET,^59^ and each sequence was simulated with explicit fluorophores using both optimized parameter sets. The original force field already captured the overall sequence dependence but yielded several chains that were too compact. The deviation from experiment correlated with the fraction of charged residues and even more strongly with the number of possible salt bridges.^34^

Because all 16 IDRs have identical length but diverse composition and charge patterning, the set provides a sensitive test of sequence-dependent intrachain interactions, while explicit treatment of the fluorophores retains their steric and electrostatic contributions. The results (Fig. 3) show that both optimized parameter sets improved the agreement with experiment relative to the original force field. The concordance correlation coefficients were 0.84 for Amber ff99SBws, 0.85 for Amber ff99SBws-CS and 0.90 for Amber ff99SBws-LJ, with the largest reductions in error for the LJ parameter set occurring among charge-rich sequences. For the subsequent, substantially more computationally demanding tests on a folded–disordered complex and protein condensates, we focused on Amber ff99SBws-LJ.

**Figure 3:**
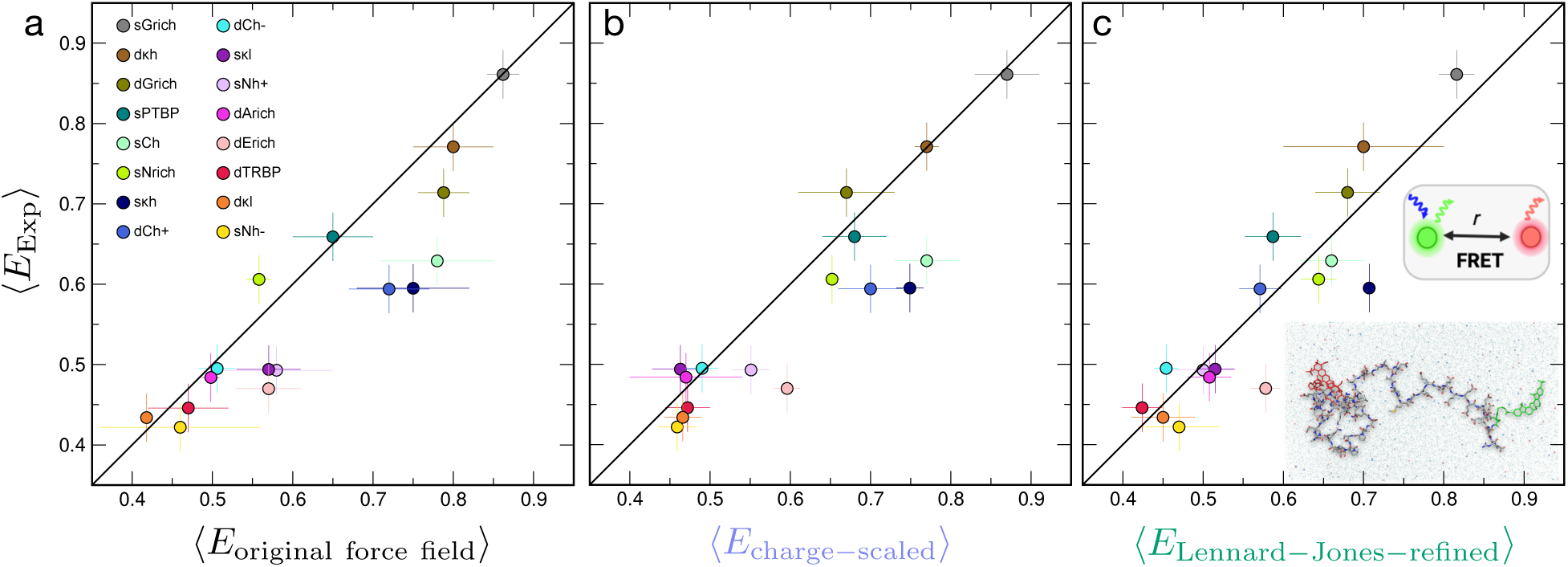
Transferability across sequence-diverse disordered proteins. Experimental mean FRET efficiencies for 16 IDRs of identical length are compared with values calculated from explicit-dye simulations using **a,** the original Amber ff99SBws force field, **b,** the charge-scaled parameters and **c,** the Lennard-Jones-refined parameters. Each color denotes one IDR (see legend). The concordance correlation coefficients are 0.84, 0.85 and 0.90, respectively. The inset shows an IDR with Cy3B (green) and CF660R (red), K^+^ (blue), Cl*^−^* (red) and water (transparent light blue); hydrogen atoms are omitted. Experimental and simulation uncertainties are defined in Methods and Refs. 34,59. Simulation error bars show the standard deviation across three independent simulations for each sequence; the corresponding standard error of the mean is 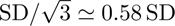. The sequences of the 16 IDRs are given in Table S1.

#### Dynamics from NMR relaxation in a folded–disordered protein complex

We next examined the highly charged complex between the intrinsically disordered prothymosin *α* (ProT*α*) and the folded globular domain of histone H1,^3^ whose structural and dynamic properties have been investigated in detail with NMR.^60^ ProT*α* remains disordered in this high-affinity complex and rapidly exchanges among many configurations on the surface of its folded partner. The system combines electrostatic interactions, structural heterogeneity and a globular protein surface, features absent from both the osmometry solutions and the monomeric-IDR validation. It is also a stringent dynamic benchmark: over-stabilized intermolecular salt bridges can slow the exchange among configurations.^60^

NMR longitudinal and transverse relaxation rates (*R*_1_ and *R*_2_) and the steady-state heteronuclear NOE report on residue-specific backbone dynamics of ProT*α*. With the original force field, simulated *R*_2_ rates for ProT*α* were substantially above the experimental values, especially in its highly charged C-terminal region (Fig. 4a). The excess *R*_2_ corresponds to motion that is too slow on the nanosecond timescale and is consistent with long-lived intermolecular contacts. Amber ff99SBws-LJ improved both *R*_1_ and *R*_2_ across the ProT*α* sequence (Fig. 4a). Intermolecular contact lifetimes provide a molecular explanation: the mean lifetime of ProT*α*–H1 contacts decreased from 12.0 ± 1.0 ns with the original parameters to 6.3 ± 0.4 ns after optimization, and the long tail of contacts persisting for hundreds of nanoseconds was strongly suppressed (Fig. 4b). Overall, the parameters optimized using amino acid osmometry improve charge-driven interactions between a disordered chain and a folded protein.

**Figure 4:**
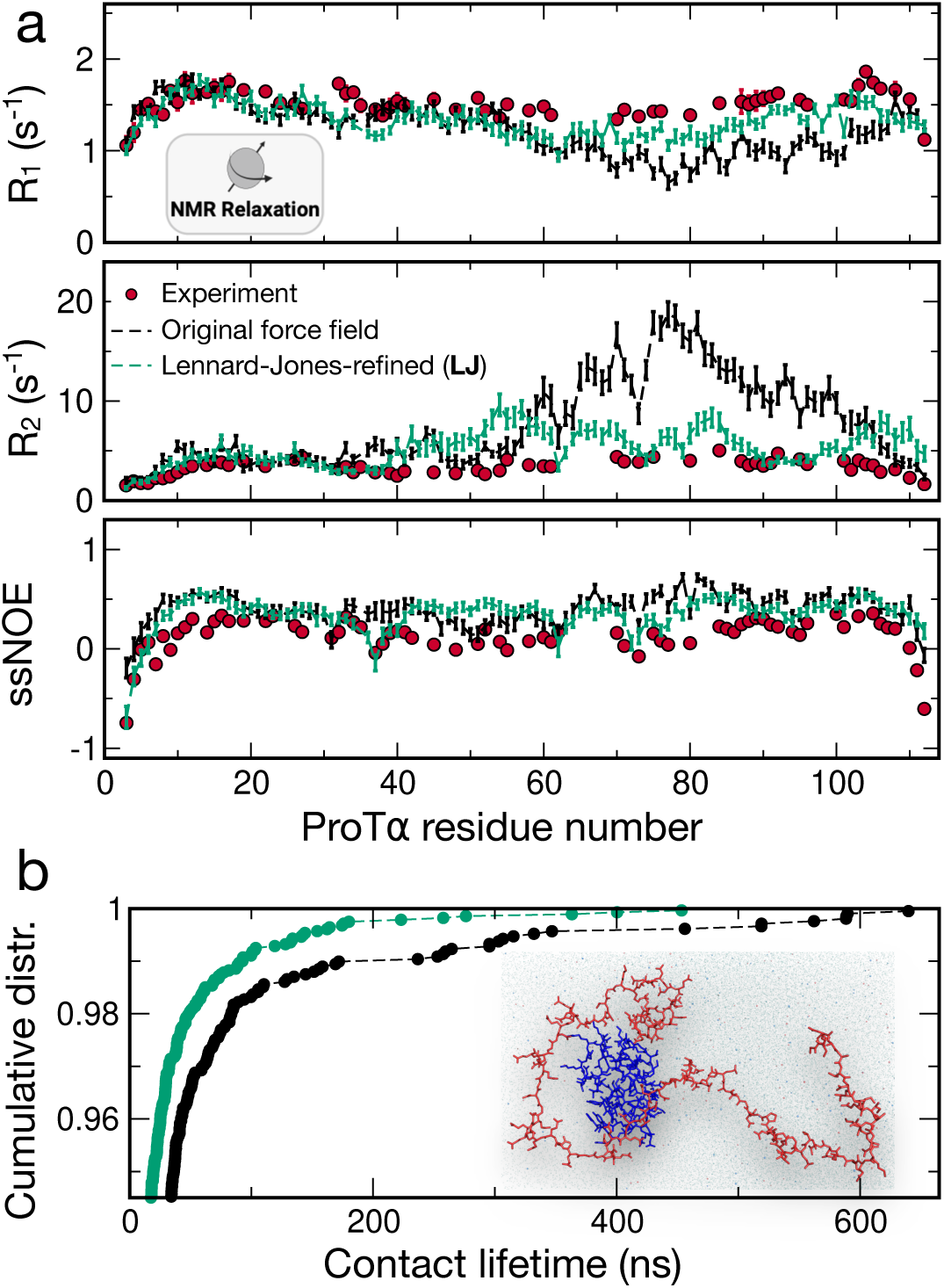
Dynamics of a complex between a folded domain and an IDP. **a,** Experimental residue-resolved *R*_1_, *R*_2_ and steady-state heteronuclear NOE values from NMR for ProT*α* bound to the globular domain of histone H1, ^60^ compared with values calculated using the original and Lennard-Jones-refined force fields. **b,** Cumulative distributions of intermolecular contact lifetimes. The optimized parameters suppress the tail of long-lived contacts present with the original force field. The inset shows ProT*α* (red), the H1 globular domain (blue), K^+^ (blue spheres), Cl*^−^* (red spheres) and water (transparent light blue); hydrogen atoms are omitted.

#### Optimized interactions match experimental chain dynamics within condensates

Multicomponent condensates provide a demanding parameter transfer test because small interaction errors can accumulate across dense networks of residue–residue and residue–ion contacts. In our previous all-atom simulations of ProT*α* condensates, the qualitative slowdown from lysine-rich to arginine-rich systems was reproduced, but ProT*α* reconfiguration in the arginine-rich condensates was 3-to 4-fold slower than in experiment.^15,16^ These systems therefore directly test whether the osmometry-optimized parameters correct a known force-field deficiency in large biomolecular assemblies. We simulated complex coacervates of ProT*α* formed either with the lysine-rich histone H1 or the arginine-rich protamine at 8 mM and 128 mM KCl, yielding four systems of 2.6–4.0 million atoms. The two polycations provide a chemically informative comparison because arginine required a larger osmometry-derived correction than lysine. The two different salt concentrations are a further test of the balance of protein and ion interactions.

We first examined ProT*α* chain dimensions within the condensates using the mean FRET efficiency, ⟨*E*⟩, for direct comparison with experiment. Amber ff99SBws already reproduces the experimental transfer efficiencies for three of the four condensate conditions, with the remaining ProT*α*–protamine condensate at low salt showing a modest deviation. Amber ff99SBws-LJ preserves the agreement across all systems and improves the agreement for the ProT*α*–protamine condensate at low salt (Table S2).

The average ProT*α* reconfiguration times in the lysine-rich ProT*α*–H1 condensates were already close to experiment with the original force field. However, a subset of chains exhibited restricted motion, defined as a standard deviation of transfer efficiency below 0.05 over the course of the simulation, suggesting trapping within a rugged energy landscape. At 128 mM KCl, 23 of 96 ProT*α* chains exhibited restricted motion with Amber ff99SBws, which was reduced to 10 with Amber ff99SBws-LJ, while the overall reconfiguration time remained similar (Fig. 5). The Lennard-Jones modifications therefore reduce local kinetic trapping without affecting the agreement already obtained for the lysine-rich condensates.

**Figure 5:**
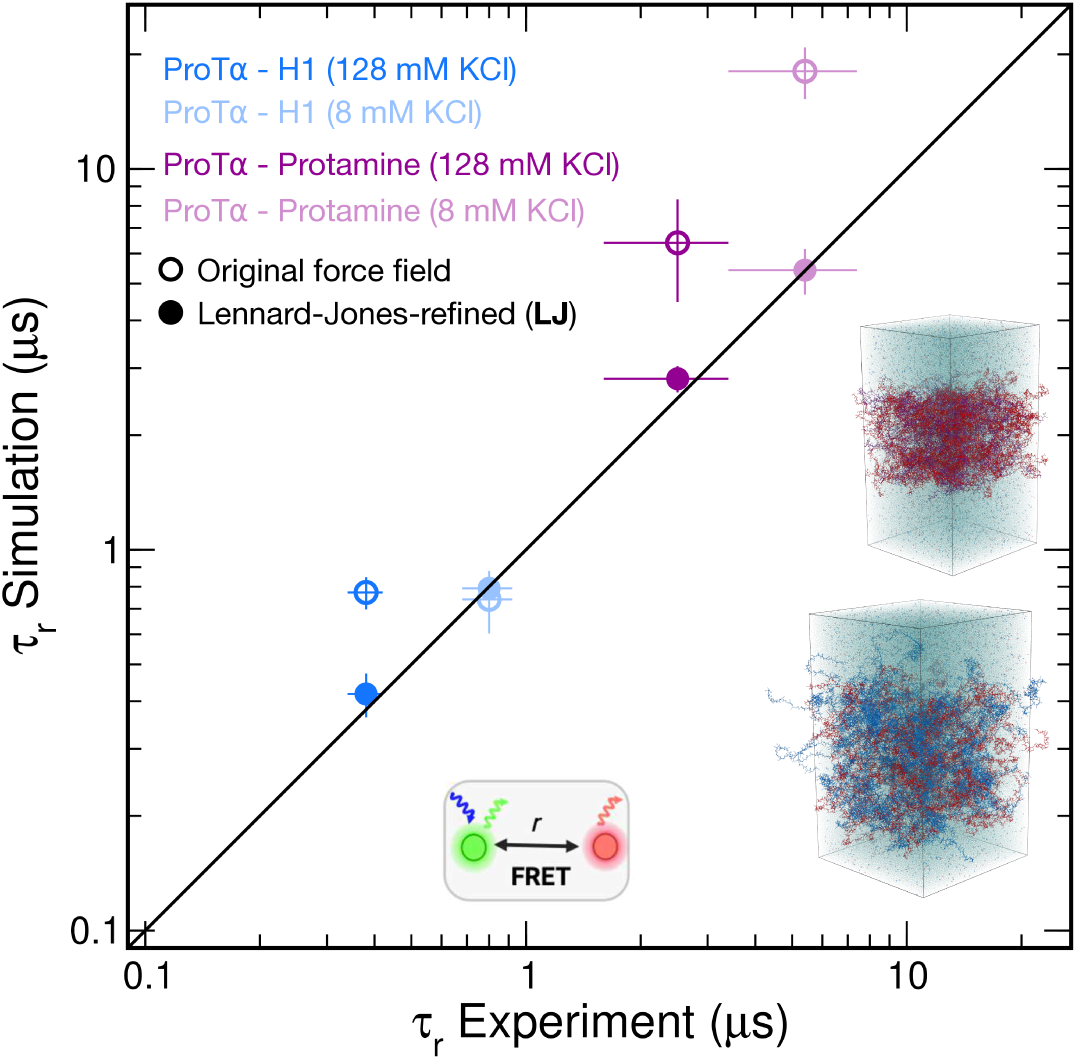
Absolute chain reconfiguration times in multicomponent condensates. ProT*α* chain reconfiguration times measured by nsFCS are compared with values calculated from all-atom simulations of condensates formed with lysine-rich H1 or arginine-rich protamine at 8 and 128 mM KCl. In the simulations, reconfiguration times were calculated from the distance between ProT*α* residues 58 and 112, corresponding to the dye positions in the experiments; the exact procedure is described in Methods. For the low-salt ProT*α*–protamine condition, the simulation at 8 mM KCl is compared directly with the closest experimental measurement, obtained at 25 mM added KCl (approximately 33 mM total ionic strength), as done previously, ^16^ because nsFCS measurements at lower salt concentrations were not feasible. Numerical values and uncertainties are given in Table S2. Open and filled circles denote the original and Lennard-Jones-refined force fields, respectively; the diagonal denotes identity. Insets show simulation snapshots of the ProT*α*–H1 and ProT*α*–protamine condensates at 128 mM KCl. ProT*α* is red, H1 blue, protamine purple, K^+^ blue, Cl*^−^* red and water transparent light blue; hydrogen atoms are omitted. Sequences of the simulated proteins are given in Table S1.

Simulations with Amber ff99SBws showed that, across the four simulated condensates, the mean lifetime of intermolecular residue–residue contacts involving ProT*α* scales with the ProT*α* reconfiguration time.^16^ We find a similar (even stronger) correlation between contact lifetimes and reconfiguration times with Amber ff99SBws-LJ (Fig. S8). The correlation in both cases strongly suggests that transient residue–residue interactions are a key determinant of chain reconfiguration, through their contribution to the effective friction for chain motion.^16^ Quantitatively reproducing reconfiguration times therefore requires an accurate balance of residue–residue interactions: by reducing overly strong salt bridges through osmometry-guided refinement, Amber ff99SBws-LJ shortens contact lifetimes (Fig. S8) and brings chain reconfiguration into better agreement with experiment (Fig. 5). The effect of the refinement is largest in the arginine-rich ProT*α*–protamine condensates. With Amber ff99SBws, ProT*α* reconfigured 3- to 4-fold more slowly than in experiment, although the qualitative slowdown relative to the ProT*α*–H1 condensates was reproduced.^16^ In simulations with Amber ff99SBws-LJ, both protamine-containing systems are in much better agreement with the experimental values (Fig. 5). The largest improvement thus occurs in the condensates enriched in arginine, the residue requiring the largest correction in the osmometry-based optimization.

Finally, we tested whether the Lennard-Jones refinement preserves the stability of folded structure. To this end, we monitored the backbone RMSD of each of the 80 globular domains of H1 relative to the NMR structure^61^ (PDB 6HQ1) over the trajectory of the ProT*α*–H1 condensate at 128 mM KCl; only 4 of 80 domains (5%) had a time-averaged RMSD above 0.4 nm (Fig. S9a). This fraction is in line with our previous simulations using Amber ff99SBws^15^ and with the experimentally determined conformational stability of the isolated GD,^61^ showing that Amber ff99SBws-LJ retains experimentally consistent folded-domain stability. Altogether, we thus arrive at a force field with an optimized representation of charge interactions, as demonstrated across a wide range of biomolecular systems, without compromising other key properties of the model, such as the stability of folded structure.

## Discussion

A central feature of our force field optimization approach is the separation of interaction-specific calibration from validation in complex biomolecular systems. Rather than optimizing parameters in proteins or condensates, where many interactions are coupled, we isolate residue–residue, residue–ion and ion–ion interactions in chemically defined solutions that retain the relevant molecular groups, but are small enough for iterative parameter refinement. Experiment and simulation are compared through the concentration dependence of the same thermodynamic observable, and only once the parameters are fixed are they transferred without further adjustment to IDPs, protein complexes and condensates.

We found that the osmometry-guided optimization yielded improvements in both configurational and dynamic properties across all three classes of biomolecular systems tested, with the most dramatic improvement observed for the dynamics within arginine-rich biomolecular condensates, consistent with the relatively large corrections to arginine required in the optimization. Notably, we achieved similar accuracy with respect to the osmotic-pressure data using two different strategies: charge scaling (Amber ff99SBws-CS), which modifies the net charges of residues, and an optimization that retains integral residue charges while modifying the Lennard-Jones parameters (Amber ff99SBws-LJ). Although formally distinct, the two strategies produce similar physical effects on charged-group association. Charge scaling weakens electrostatic attraction between charged groups, whereas Lennard-Jones optimization does so indirectly: reducing the residue–water distance permits water molecules to approach the charged groups more closely, thereby increasing solvation of the oppositely charged groups.

A driving force for recent force field improvements has been variational optimization against experimental data reflecting protein conformations in solution.^19,21,24,25^ The osmometry-guided strategy developed here provides a different source of experimental information: chemically defined solutions isolate specific intermolecular interaction classes, and their measured thermodynamics provide direct targets for parameter optimization. The value of optimized force fields with the accuracy we achieve here is illustrated by recent developments that link molecular-scale dynamics to the mesoscopic properties of biomolecular condensates. For complex coacervates of the type we simulated here (Fig. 5), relations from polymer physics can be used to predict the bulk viscosity and translational diffusion coefficients of the constituent IDPs from their chain reconfiguration times.^16^ If the reconfiguration time can be obtained directly from simulations with sufficient accuracy, it is thus possible to predict mesoscopic properties of such systems from simulations alone.

The role of charge interactions extends far beyond protein salt bridges: they contribute to protein–DNA association, complexation of arginine-rich RNA-binding regions with RNA, and the assembly and selective composition of transcriptional, nucleolar and stress-associated condensates.^6,62,63^ More generally, the strategy developed here can be used when new forcefield deficiencies arise in increasingly complex biomolecular simulations. Chemically defined osmometry measurements can be designed to separate candidate interaction classes, identify which interactions are responsible for a discrepancy, and provide a direct experimental target for their optimization before the optimized parameters are transferred back to the complete biomolecular system. Protein–nucleic-acid interactions are a promising next application of this strategy, for instance the interactions of phosphate groups with basic protein side chains and ions, since conventional additive force fields can over-stabilize amine–phosphate interactions in peptide–nucleic-acid assemblies.^64^ In this work, we used osmometry-guided optimization to correct deficiencies identified in an existing force field. Because the approach is computationally inexpensive, it could also be incorporated directly into de novo force-field development as part of a unified optimization scheme.

## Methods

### Osmometry measurements

L-Lysine monohydrate, L-arginine, L-glutamic acid, L-aspartic acid, L-aspartic acid potassium salt and L-aspartic acid sodium salt monohydrate were purchased from Sigma-Aldrich. Equimolar mixtures of oppositely charged amino acids (Lys–Glu, Lys–Asp, Arg–Glu and Arg–Asp) were prepared by combining L-lysine monohydrate or L-arginine with L-glutamic acid or L-aspartic acid. The amounts of the basic and acidic amino acids were determined by weighing each powder with a precision of 0.0003 g. The highest molality prepared for each amino acid pair was approximately 0.55 mol kg*^−^*^1^, with the exact value chosen to match the molality of the corresponding simulated system. Solutions of lower molality were prepared by dilution with ultrapure water. The water of crystallization in L-lysine monohydrate and L-aspartic acid sodium salt monohydrate was included when calculating the sample concentrations. Solutions containing Asp–Na^+^ and Asp–K^+^ were prepared by dissolving the corresponding purchased salts and subsequently diluting the resulting solutions. All osmometry measurements were performed using a VAPRO Vapor Pressure Osmometer 5600. Calibration was performed at the beginning of each measurement day using 100 mOsm kg*^−^*^1^, 290 mOsm kg*^−^*^1^ and 1000 mOsm kg*^−^*^1^ calibration standards. The standards were then remeasured until all three gave stable readings within 3 mOsm kg*^−^*^1^ of their nominal values.

Measured osmolality values were converted to osmotic pressure using Π = *c*_osm_*RT*, where *c*_osm_ is the concentration of osmotically active particles corresponding to the measured osmolality, *R* is the gas constant and *T* = 298.15 K. For each of the six datasets shown in Fig. 2 (Lys–Glu, Lys–Asp, Arg–Glu, Arg–Asp, Asp–Na^+^ and Asp–K^+^), samples were prepared using at least two independently purchased batches, and every concentration point was measured on at least two different days. At each concentration on each measurement day, three to six replicate measurements were performed and averaged to obtain a daily mean. The reported osmotic pressure for each concentration point was calculated as the mean of the daily means. Uncertainties for all reported osmotic-pressure values were estimated by combining in quadrature the manufacturer-specified instrumental uncertainty of 1% of the reported osmotic pressure and a calibration contribution corresponding to the 3 mOsm kg*^−^*^1^ calibration tolerance. At 298.15 K, the latter corresponds to 0.0744 bar, giving 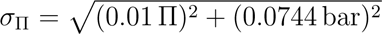, where Π is the reported osmotic pressure. For concentration points measured independently on at least three days, the standard deviations among the daily means were also calculated, which were similar to the corresponding values of *σ*_Π_ calculated above. The value of *σ*_Π_ calculated as shown above was therefore used as the uncertainty estimate for every reported osmotic-pressure value.

### Derivation of force-field parameters for amino acids with charged termini

Since the Amber force fields do not include standard parameters for single residues in which both termini are charged (i.e., zwitterions, in the case of a neutral side-chain), it was necessary to determine such parameters. We used the Amber Antechamber tool to automatically assign protein atom types and bonded parameters (angles, dihedrals) involving the terminal atoms, keeping the bonded parameters for the side-chain the same as in the protein force field. This left the atomic partial charges as the remaining critical parameter. For Amber force fields, partial charges are usually fitted to a molecular electrostatic potential via the RESP algorithm. However, we made several assumptions to reduce the degeneracy of the problem and to maximize similarity with the protein force field. First, we fixed the sidechain charges (i.e. those on the *β* position of the side-chain and beyond) to be identical to those for each residue/amino acid in the standard protein force field, with the aim of improving transferability of the results. Second, we assumed that a common set of backbone charges should be used across all residues. This was motivated by the Cornell *et al.* force field (Amber 94)^65^ and its many descendants using the same charges on backbone atoms for all internal residues, except for the residues with charged side-chains. We had subsequently found that using common charges for all residues (including charged residues) in fact resulted in improved prediction of *α*-helix propensity,^66^ and this change was incorporated in the Amber ff99SBws.^20^ Recent electron-diffraction measurements of zwitterionic tyrosine and histidine provide independent experimental support for this approximation: the summed partial charges of the NH_3_, C*_α_*H*_α_* and COO groups are similar for the two amino acids.^67^ Therefore, the problem reduces to finding a set of charge differences (summing to zero) from the fixed charge set for internal residues. We determined these differences by (i) running a standard RESP fit^68^ of charges to an electrostatic potential computed with the HF/6-31G basis set using Gaussian 16;^69^ (ii) computing differences of the backbone charges from those for interior residues for each of lysine, arginine, glutamate and aspartate; (iii) averaging the differences and subtracting their sum (0.0259) from the *α* carbon charge difference to give a net difference of zero; (iv) adding these differences to the standard force-field charges. This scheme preserves the total charge of each residue. We applied the same backbone charges also to form the zwitterions of alanine and glycine.

### All-atom osmometry simulations

Osmotic pressures were calculated from all-atom simulations using a virtual-semipermeable-wall approach based on the method introduced by Luo and Roux.^40^ In their original implementation, Luo and Roux calculated osmotic pressures for concentrated NaCl and KCl aqueous solutions by confining the ions to a defined region of the simulation box with virtual walls, while allowing water molecules to move freely through the periodic system. The osmotic pressure was then obtained from the mean force exerted by the virtual walls on the confined solutes, divided by the wall area. Here, the virtual walls were implemented in GROMACS^70^ using flat-bottomed position restraints. Water molecules were not restrained and could diffuse freely throughout the simulation box. The solute particles used to calculate the osmotic pressure were free to diffuse in the *x* and *y* directions and within the defined slab along the *z* direction. When a restrained solute particle moved outside this slab, the flat-bottomed restraint applied a harmonic restoring force directed back toward the confined region. Thus, the restraints acted as idealized semipermeable walls that confined the solutes but not the water. The flat-bottomed restraints were applied with a layer geometry normal to the *z* direction. For atomic ions, the restraint was applied directly to the ion. For amino acid residues, the restraint was applied to the C*_α_* atom of each residue. The same restraint parameters were assigned to all residues of a given type through the corresponding residue topology. The osmotic pressure was calculated from the force exerted by the virtual walls on the restrained solute particles, following Luo and Roux.^40^ At each saved simulation frame, the force contribution from each restrained particle outside the flat-bottom region was calculated from the applied restraint. The force magnitudes from the two slab boundaries were averaged, and the osmotic pressure was calculated as

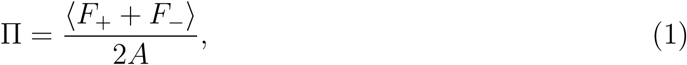

where *F*_+_ and *F_−_* are the instantaneous force magnitudes exerted by the left and right virtual walls, respectively, and *A* is the cross-sectional area of the simulation box perpendicular to the *z* direction. This gives the osmotic pressure directly from the mechanical force required to confine the solutes while allowing water to equilibrate across the virtual semipermeable boundary.

To validate the GROMACS implementation, we repeated the NaCl and KCl osmometry simulations reported by Luo and Roux.^40^ The validation simulations used the same forcefield description as their final optimized simulations, namely CHARMM ion parameters together with TIP3P water as modified for the CHARMM force field, including the optimized cation–anion NBFIX parameters introduced in their work. Across the full NaCl and KCl concentration ranges reported by Luo and Roux, the osmotic pressures obtained with our flatbottom-restraint implementation reproduced their values, confirming that the GROMACS setup gives the same osmotic pressure as the original virtual-wall approach. For the NaCl and KCl simulations, rectangular boxes were used, with dimensions of 5 nm in the *x* and *y* directions and 10 nm in the *z* direction. Ions were confined to the central 5 nm region along the *z* axis using the flat-bottomed restraints described above. For both NaCl and KCl, we simulated nominal molar concentrations of 0.1 M, 0.2 M, 0.3 M, 0.4 M, 0.5 M, 1 M, 2 M, 3 M, and 4 M, with an additional 5 M simulation for NaCl. The exact concentrations differed slightly from the nominal values because the systems contained integer numbers of ions. The corresponding molal concentrations were determined from the number of ions and the average number of water molecules between the virtual walls,^41,42^ enabling direct comparison with osmotic pressures reported as a function of molal salt concentration.^46^ For each NaCl and KCl system, a 5 ns NPT simulation was first performed using semi-isotropic pressure coupling with the Parrinello–Rahman barostat^71^ at a reference pressure of 1 bar. Only the *z* dimension of the box was allowed to fluctuate, with *τ_p_* = 5 ps. The temperature was maintained at 298.15 K using the velocity-rescaling thermostat.^72^ Long-range electrostatic interactions were treated using the particle-mesh Ewald method.^73^ Dispersion interactions and short-range repulsion were described by a Lennard-Jones potential with a cutoff of 1 nm. Hydrogen-bond lengths were constrained using the LINCS algorithm.^74^ A 2 fs integration time step was used. The final structures from the NPT simulations were used as starting structures for production simulations. Production simulations were performed in the NVT ensemble to keep the box volume fixed during osmotic-pressure calculations. All simulations used for the final reported osmotic pressures were 502 ns long. Shorter simulations were used only during parameter testing, when the osmotic pressure already showed a clear deviation from experiment; in these cases, simulations were stopped after at least 100 ns to avoid spending additional computational resources on parameter sets that were already inconsistent with the experimental data. Ion coordinates were saved every 1 ps, and the first 2 ns of each production simulation were discarded as equilibration. Osmotic pressures were calculated from the ion coordinates at each saved frame. For the final reported 502 ns simulations, the reported values were obtained from the final 500 ns of each trajectory, and uncertainties were estimated as the standard deviation of the mean osmotic pressures calculated from five consecutive 100 ns blocks. For the shorter parameter-testing simulations, the same block-averaging procedure was applied to the available production trajectory after discarding the initial 2 ns equilibration period. We note that we were able to obtain equivalent osmotic pressure values by computing the virial pressure from the total forces (including solvent) acting on the ions, although with greater statistical uncertainty.

Residue–ion simulations were performed using the same virtual-semipermeable-wall setup as the NaCl and KCl simulations. The simulated residue–ion systems were Lys–Cl*^−^*, Arg– Cl*^−^*, Glu–K^+^, Glu–Na^+^, Asp–K^+^, and Asp–Na^+^. For these systems, cubic boxes were used, with dimensions of 10 nm along each of the three axes. Residues and ions were confined to the central ∼5 nm region of the box along the *z* direction. The exact width of the restrained region along *z* was chosen such that the nominal molar concentrations of each residue–ion system were 100 mM, 250 mM, 350 mM, and 500 mM. Throughout the manuscript, osmotic pressures are reported as a function of molal concentration. The molal concentration was determined from the number of solute particles and the average number of water molecules in the central restrained region during the production simulations. For each residue–ion system and concentration, a 25 ns NPT simulation was first performed. From each NPT simulation, five equally spaced snapshots were extracted every 5 ns and used as starting structures for five independent production simulations. Each production simulation was 252 ns long, with the first 2 ns discarded as equilibration. All other simulation parameters were identical to those used for the NaCl and KCl simulations. Osmotic pressures were calculated every 1 ps from the restrained-particle coordinates. For each residue–ion system and concentration, the reported osmotic pressure was calculated as the mean over the five independent production simulations. Uncertainties were reported as the standard deviation across the mean osmotic pressures obtained from the five independent simulations.

Residue–residue simulations of oppositely charged pairs (Lys–Glu, Lys–Asp, Arg–Glu, and Arg–Asp) were performed using the same simulation protocol as the residue–ion systems. The only difference was the set of simulated concentrations: for each residue–residue pair, nominal molar concentrations of 200 mM, 300 mM, 400 mM, and 500 mM were used. As for the residue–ion simulations, molal concentrations were determined from the number of solute particles and the average number of water molecules in the central restrained region during the production simulations. Single-residue simulations of alanine and glycine were performed using the same setup and analysis procedure. For these systems, nominal molar concentrations of 100 mM, 250 mM, 500 mM, 1 M, and 2 M were simulated. For each residue– residue and single-residue system, a 25 ns NPT simulation was first performed, followed by five independent 252 ns production simulations initiated from five equally spaced snapshots extracted every 5 ns from the NPT trajectory. The first 2 ns of each production simulation were discarded as equilibration, and osmotic pressures were calculated every 1 ps from the restrained-particle coordinates. Reported osmotic pressures were calculated as the mean over the five independent production simulations, and uncertainties were reported as the standard deviation across the mean osmotic pressures obtained from these five simulations.

To calculate coordination numbers at near-saturation salt concentrations, we performed simulations at 6.18 mol kg*^−^*^1^ for NaCl and 4.56 mol kg*^−^*^1^ for KCl.^57^ Simulations were performed in cubic boxes with a side length of 7 nm. The numbers of Na^+^, K^+^, and Cl*^−^* ions were chosen such that, after replacing water molecules with ions, the target experimental molal concentrations were obtained.^57^ For both NaCl and KCl, simulations were performed for the initial and all relevant modified force-field parameter sets. All simulations were at least 25 ns long, and the first 1 ns was discarded as equilibration. All other simulation parameters were identical to those used for the ion–ion osmometry simulations. Following previous work,^75^ coordination numbers were determined from the Na^+^–Cl*^−^* and K^+^–Cl*^−^* radial distribution functions. The radial distribution functions were calculated using a bin width of 0.001 nm. The position of the first minimum was determined by fitting the RDF in a narrow region around the local minimum to a quadratic function, *f* (*r*) = *ar*^2^ + *br* + *c*, with the minimum position calculated as *r*_min_ = −*b/*(2*a*). Coordination numbers were then calculated by integrating the RDF up to this first minimum, 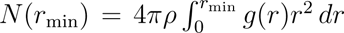, where *ρ* is the bulk number density of the counterion species, *g*(*r*) is the ion–ion radial distribution function, and *r*_min_ is the position of the first minimum. Uncertainties were estimated using block analysis.

#### All-atom simulations of IDRs

All-atom simulations of the 16 dye-labelled IDRs were performed using the same solvated starting structures as in our previous work.^34^ For each IDR, two sets of simulations were carried out using the osmometry-guided force-field modifications described in the main text: one using the optimized charge-scaling parameters and one using the optimized Lennard-Jones *σ* parameters. The previously prepared dye-labelled systems, including water and ions, were reused directly; only the force-field parameters were changed before rerunning the simulations. The starting systems contained all-atom configurations of the 16 IDR variants labelled with Cy3B and CF660R in 14 nm rhombic dodecahedral boxes with TIP4P/2005s water.^20^ Potassium and chloride ions were present at 172 mM KCl, matching the total ionic strength of the KCl salt and buffer used in the experiments. The total number of atoms per simulation was approximately 250,000, varying slightly among different IDRs. All simulations were performed with GROMACS version 2021.5.^70^ Protein interactions were modeled using Amber ff99SBws,^20^ together with the osmometry-guided parameter modifications described in the main text. Cy3B and CF660R were described using previously published dye parameters, and transfer efficiencies were calculated using the Förster radius of *R*_0_ = 6.0 nm.^76,77^ Dye–water Lennard-Jones interactions were scaled by a factor of 1.15 for both Cy3B and CF660R, as described previously.^78^

For simulations with the optimized charge-scaling parameters, the charges of the charged groups within the dyes were scaled by the same factor as the charged protein residues, 0.93. For simulations with the optimized Lennard-Jones parameters, the Lennard-Jones *σ* parameters between the charged dye groups and water oxygen atoms were scaled by a factor of 0.99. This factor was chosen to match the average magnitude of the residue-specific *σ* modifications applied to charged residue–water oxygen interactions: *σ*_Lys–OW_ and *σ*_Asp–OW_ were scaled by 0.99, *σ*_Glu–OW_ by 0.995, and *σ*_Arg–OW_ by 0.985. The integration time step was 2 fs. The temperature was maintained at 295.15 K using the velocity-rescaling thermostat^72^ with *τ_t_* = 1 *ps*, and the pressure was maintained at 1 bar using the Parrinello– Rahman barostat^71^ with *τ_p_* = 5 *ps*. Long-range electrostatic interactions were treated using the particle-mesh Ewald method.^73^ Dispersion interactions and short-range repulsion were described by a Lennard-Jones potential with a cutoff of 1 nm. Hydrogen-bond lengths were constrained using the LINCS algorithm.^74^ Each independent simulation was at least 2.02 *µ*s long, and the first 20 ns were discarded as equilibration.

#### All-atom simulations of folded–disordered complex dynamics

Simulations of the complex between the globular domain (GD) of histone H1 and ProT*α* were run using a protocol similar to that used previously.^60^ The initial protein configuration was taken from the earlier simulations and solvated with explicit water and 165 mM potassium chloride in a 17 nm truncated-octahedral box with periodic boundary conditions. Either the original Amber ff99SBws model or the Amber ff99SBws-LJ model was used for the protein and ions, with TIP4P/2005s water.^50^ The total number of atoms was 495,529. Simulations were performed using GROMACS 2024.1,^70^ with the equations of motion integrated using the velocity-Verlet algorithm with a time step of 2 fs and LINCS constraints on all bonds. A constant temperature of 283 K was maintained using the Bussi velocity rescaling thermostat^72^ with a 1 ps relaxation time, and a constant pressure of 1 atm was maintained using the Parrinello–Rahman barostat^71^ with a 5 ps relaxation time. Simulations were run for 5 *µ*s in total, with the first 3 *µ*s discarded as equilibration. NMR relaxation parameters were computed as previously described.^60^

Residue–residue contact lifetimes were calculated using a transition-based core-state approach,^79^ as described previously.^15,16^ In short, for each pair of residues belonging to different protein molecules, the shortest distance between any pair of heavy atoms was monitored. Contact formation was defined by this distance decreasing below 0.38 nm, whereas an existing contact was considered broken only when the distance increased above 0.8 nm. The mean lifetime of a residue–residue contact was calculated by dividing its total bound time by the number of contact-breaking events.

#### All-atom simulations of biomolecular condensates

All-atom simulations of ProT*α*–H1 and ProT*α*–protamine condensates at 8 mM and 128 mM KCl were performed using the same system compositions and simulation protocols as in our previous studies,^15,16^ with the force-field parameters changed as described below. The previous simulations used Amber ff99SBws, referred to as the “original force field” in the present manuscript, whereas all new simulations used Amber ff99SBws-LJ. The ProT*α*–H1 simulation at 128 mM KCl was initiated from the snapshot at 4 *µ*s of the corresponding previously reported Amber ff99SBws trajectory.^15^ A snapshot at 1.45 *µ*s of the resulting Amber ff99SBws-LJ trajectory was used to initiate the ProT*α*–H1 simulation at 8 mM KCl. The ProT*α*–protamine simulation at 128 mM KCl was initiated from the same starting structure as the corresponding previously reported Amber ff99SBws simulation,^16^ and a snapshot at 3.45 *µ*s of the resulting Amber ff99SBws-LJ trajectory was used to initiate the ProT*α*– protamine simulation at 8 mM KCl.

All systems were simulated under periodic boundary conditions in slab geometry with TIP4P/2005s water.^50^ The temperature was maintained at 295.15 K using stochastic velocity rescaling^72^ with a relaxation time of 1 ps, and the pressure was maintained at 1 bar using the Parrinello–Rahman barostat.^71^ Long-range electrostatic interactions were treated using the particle-mesh Ewald method^73^ with a grid spacing of 0.12 nm. Lennard-Jones interactions were truncated at 0.9 nm, bonds involving hydrogen atoms were constrained using LINCS, and the equations of motion were integrated using the leap-frog algorithm with a time step of 2 fs. Simulations were performed using GROMACS version 2024.1.^70^

The ProT*α*–H1 simulations contained 96 ProT*α* and 80 H1 molecules. The system at 128 mM KCl contained 4,000,932 atoms and was simulated for 5.5 *µ*s, with the first 1.5 *µ*s excluded as equilibration. The system at 8 mM KCl contained 3,996,354 atoms and was simulated for 6.8 *µ*s, with the first 1.8 *µ*s excluded as equilibration. The ProT*α*–protamine simulations contained 96 ProT*α* and 197 protamine molecules. The system at 128 mM KCl contained 2,612,851 atoms and was simulated for 10.0 *µ*s, with the first 2.2 *µ*s excluded as equilibration. The system at 8 mM KCl contained 2,609,869 atoms and was simulated for 8.4 *µ*s, with the first 1.0 *µ*s excluded as equilibration. Equilibration was assessed from equilibration of the protein density in the central region of the condensate slab, as described previously (Fig. S9).^15,16^

The stability of the H1 globular domains (Fig. S9a) in the ProT*α*–H1 condensate at 128 mM KCl was assessed as previously described.^15^ Backbone RMSDs were calculated for each of the 80 GDs relative to the experimental structure (PDB 6HQ1)^61^ using snapshots sampled every 50 ns. Domains with a mean RMSD above 0.4 nm were classified as partially unfolded, following the criterion used previously.^15^

Protein mass concentrations in the dense phases (Fig. S9b) were calculated from protein mass-density profiles along the *z* axis, as described previously.^15,16^ The trajectories were divided into 50 ns blocks, and a protein mass-density profile was calculated for each block. The dense-phase concentration was obtained by averaging the profile over the central region of the condensate slab, excluding the interfacial regions. The boundaries of the central region were adjusted over the trajectory to account for changes in the position and width of the slab. For ProT*α*–H1 at 128 mM KCl, the following central regions were used: 21.0 *< z <* 37.1 nm from 0 to 0.50 *µ*s, 20.0 *< z <* 38.1 nm from 0.50 to 1.00 *µ*s, 19.5 *< z <* 37.1 nm from 1.00 to 1.50 *µ*s, 18.8 *< z <* 36.6 nm from 1.50 to 2.10 *µ*s, 18.8 *< z <* 36.8 nm from 2.10 to 2.30 *µ*s, 18.8 *< z <* 37.0 nm from 2.30 to 2.45 *µ*s, 19.0 *< z <* 37.0 nm from 2.45 to 2.85 *µ*s, 18.0 *< z <* 36.0 nm from 2.85 to 3.05 *µ*s, 18.3 *< z <* 36.3 nm from 3.05 to 3.20 *µ*s, 17.5 *< z <* 37.0 nm from 3.20 to 3.50 *µ*s, 18.5 *< z <* 37.5 nm from 3.50 to 4.35 *µ*s, and 18.0 *< z <* 38.1 nm from 4.35 to 5.50 *µ*s. For ProT*α*–H1 at 8 mM KCl, the regions were 18.5 *< z <* 36.5 nm from 0 to 0.70 *µ*s, 17.5 *< z <* 36.5 nm from 0.70 to 1.40 *µ*s, 18.5 *< z <* 36.1 nm from 1.40 to 2.75 *µ*s, 19.5 *< z <* 37.1 nm from 2.75 to 4.35 *µ*s, 20.1 *< z <* 36.7 nm from 4.35 to 4.85 *µ*s, 20.1 *< z <* 36.1 nm from 4.85 to 5.50 *µ*s, and 19.0 *< z <* 36.7 nm from 5.50 to 6.80 *µ*s. For ProT*α*–protamine at 128 mM KCl, the regions were 13.0 *< z <* 30.0 nm from 0 to 0.10 *µ*s, 12.0 *< z <* 29.0 nm from 0.10 to 0.25 *µ*s, 12.0 *< z <* 28.0 nm from 0.25 to 0.60 *µ*s, 13.0 *< z <* 27.5 nm from 0.60 to 0.80 *µ*s, 12.5 *< z <* 25.5 nm from 0.80 to 1.00 *µ*s, 13.0 *< z <* 26.5 nm from 1.00 to 1.80 *µ*s, 12.7 *< z <* 26.0 nm from 1.80 to 6.00 *µ*s, 12.3 *< z <* 25.3 nm from 6.00 to 8.50 *µ*s, and 13.3 *< z <* 25.7 nm from 8.50 to 10.00 *µ*s. For ProT*α*–protamine at 8 mM KCl, the regions were 13.0 *< z <* 30.0 nm from 0 to 0.10 *µ*s, 12.0 *< z <* 29.0 nm from 0.10 to 0.25 *µ*s, 12.0 *< z <* 28.0 nm from 0.25 to 0.60 *µ*s, 13.0 *< z <* 27.5 nm from 0.60 to 0.80 *µ*s, 12.5 *< z <* 25.5 nm from 0.80 to 1.00 *µ*s, 13.0 *< z <* 26.5 nm from 1.00 to 1.80 *µ*s, 12.7 *< z <* 25.5 nm from 1.80 to 5.00 *µ*s, 12.5 *< z <* 24.5 nm from 5.00 to 6.50 *µ*s, and 13.0 *< z <* 24.1 nm from 6.50 to 8.40 *µ*s. For each simulation, the reported protein mass concentration was calculated as the arithmetic mean of the 50 ns block values within the analyzed trajectory segment indicated in Fig. S9. Its uncertainty was taken as the standard deviation of these block values.

The transfer efficiency of ProT*α* was calculated as previously described.^15,16^ In brief, the instantaneous transfer efficiency was calculated every 50 ps for each ProT*α* chain using

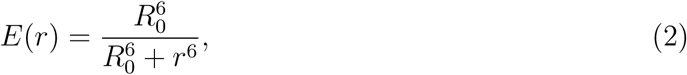

with a Förster radius of *R*_0_ = 5.9 nm, reduced from 6.0 nm owing to the increased refractive index in the condensates.^16^ Because the fluorophores and their linkers were not represented explicitly, the interdye distance was estimated from the distance, *d*, between the C*_α_* atoms of ProT*α* residues 58 and 112 according to

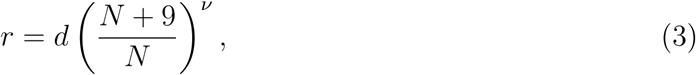

where *N* is the sequence separation between the labelling sites and *ν* = 0.6. The additional nine effective residues account for the combined lengths of the fluorophores and their linkers. For each ProT*α* chain, *E*(*r*) was averaged over the analyzed trajectory. The condensate-averaged mean transfer efficiency, ⟨*E*⟩, was calculated by averaging these chain-specific means over all 96 ProT*α* chains. The reported simulation dispersion is the standard deviation across these 96 chain-specific mean transfer efficiencies and therefore describes chain-to-chain heterogeneity rather than the uncertainty in the condensate mean.

Residue–residue contact lifetimes were calculated as for the ProT*α*-GD system described above. The analysis included all intermolecular contacts involving ProT*α*: contacts between different ProT*α* molecules and between ProT*α* and H1 or protamine. Intrachain, H1–H1 and protamine–protamine contacts were excluded. Given the large number of included contacts, each of the four simulations performed with Amber ff99SBws-LJ was divided into blocks of 1–2 *µ*s, and each block was analyzed independently using parallelized code. A mean contact lifetime was obtained for each block by averaging over all included contacts. For each simulation, the reported contact lifetime is the mean of the block-specific values, and its uncertainty is their standard deviation. The corresponding values for the four simulations performed with the original Amber ff99SBws force field were taken from our previous analyses, which used the same contact definition and lifetime calculation.^15,16^

Reconfiguration times were determined from the distance, *r*(*t*), between the C*_α_* atoms of residues 58 and 112 of each ProT*α* chain for all four simulations performed with Amber ff99SBws-LJ and the four corresponding simulations performed previously with Amber ff99SBws.^15,16^ The same one-dimensional Smoluchowski diffusion model as in our previous work was used.^16^ Briefly, the distance coordinate between 2 nm and 10 nm was discretized into 30 equal-width bins. For each candidate lag time, Δ*t*, transitions from bin *i* to bin *j* were counted separately within each ProT*α* trajectory and subsequently pooled over the 96 chains. The discretized free energies and a constant diffusion coefficient were optimized using the previously described Monte Carlo likelihood procedure,^16,34^

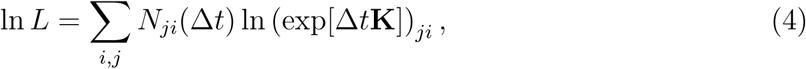

where *N_ji_*(Δ*t*) is the number of observed transitions and **K** is the rate matrix constructed from the free energies and diffusion coefficient according to the discretized Smoluchowski scheme of Bicout and Szabo.^80,81^ The reconfiguration time was calculated from the eigenvalues, *λ_n_*, and right eigenvectors, **Ψ***^R^*, of **K** as

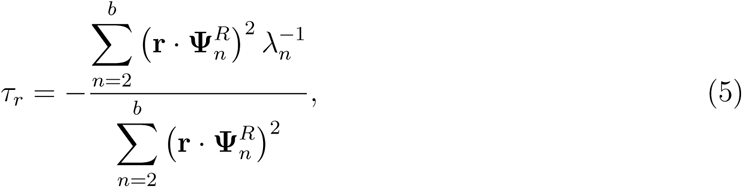

where the elements of **r** are the centers of the distance bins.

In our previous analysis,^16^ a common lag time of 200 ns was used for all systems based on the apparent convergence of *τ_r_* with lag time, and the values obtained at 100 ns and 300 ns were used to estimate the associated lag-choice uncertainty. Several of the new trajectories are substantially longer than those analyzed previously, allowing a more stringent assessment of lag dependence. We therefore reanalyzed all eight trajectories using a system-specific predictive criterion rather than imposing the same lag time on every system. The lag used to define the lag-time averaging window was selected using a Chapman–Kolmogorov (CK) predictive test.^82^ For each candidate lag, the fitted rate matrix was propagated to 2Δ*t*, 3Δ*t*, and 4Δ*t*, and the resulting model transition matrices were compared with empirical transition matrices counted directly from the trajectories at the corresponding longer lag times. At each prediction time, the discrepancy was quantified using a population-weighted total-variation distance,

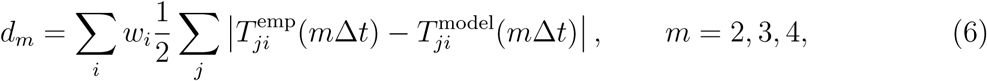

where the origin bins were weighted by their observed populations, *w_i_*.

To determine the magnitude of the discrepancy expected from finite sampling alone, complete ProT*α* trajectories were resampled with replacement. Each bootstrap replicate contained 96 trajectories drawn from the original set of 96 ProT*α* trajectories, thereby preserving the temporal correlations within each trajectory. For every bootstrap replicate, empirical transition matrices were recalculated at 2Δ*t*, 3Δ*t*, and 4Δ*t*, and their population-weighted total-variation distances from the corresponding full-data empirical matrices were determined. The 95th percentile of this bootstrap distribution, 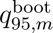, defined the finite-sampling noise threshold at each prediction time. Only prediction times retaining at least 200 estimated independent transitions were included. The CK score was calculated as

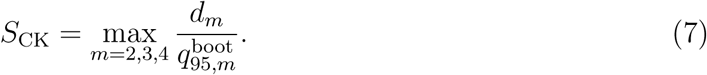

A CK-score acceptance criterion of *S*_CK_ ≤ 1 was used; values of *S*_CK_ *>* 1 were therefore classified as not passing the CK test. The smallest lag that passed the CK test was selected as the reference lag. For passing cases, the reported *τ_r_* was the unweighted mean over all CK-passing candidate lags, and the sample standard deviation of *τ_r_* across these lags was reported as a descriptive measure of its lag dependence. If no candidate lag passed the CK test, the lag with the lowest CK score was selected as the reference lag, and the result was reported as a best-effort, not-converged estimate. The averaging set in these cases comprised the selected lag and adjacent, sufficiently sampled candidate lags with CK scores no more than 25% above the minimum score. The reported *τ_r_* was again the unweighted mean over this lag set, with its sample standard deviation reported as the lag-dependence uncertainty.

As a control for the CK-based analysis, we applied the same procedure to interdyedistance trajectories from 15 independent all-atom simulations of the explicitly dye-labelled disordered peptide G(AGQ)_6_AGC, each approximately 1.1 *µ*s long.^78^ The CK test passed at a reference lag of 3.5 ns, and the CK-passing candidate lags used for averaging were 3.5, 4, 4.5, 5, 6, 7, 8, 9, 10, 12, 14, 15, 20, 25 and 50 ns, yielding a reconfiguration time of (14±2) ns. This value agrees with the previously reported values of 15 ns from direct autocorrelation of the MD interdye distance and 14 ns from a one-dimensional diffusion analysis based on simulated photon-emission data.^78^

For the ProT*α*–H1 condensate at 128 mM KCl, the CK test passed for both force fields. With Amber ff99SBws, the reference lag was 625 ns, and the CK-passing candidate lags used for averaging were 625, 750, 875, and 1000 ns. With Amber ff99SBws-LJ, the reference lag was 500 ns, and the CK-passing candidate lags used for averaging were 500, 600, 700, 800, and 900 ns.

For the ProT*α*–H1 condensate at 8 mM KCl, no candidate lag passed the CK test for Amber ff99SBws. The lowest CK score was 1.2 at a reference lag of 200 ns, and the local lag set used for averaging comprised 100 and 200 ns. For Amber ff99SBws-LJ, the CK test passed, the reference lag was 700 ns, and the CK-passing candidate lags used for averaging were 700, 800, 825, 850, 900, 950, 975, and 1000 ns. The analyzed part of the Amber ff99SBws trajectory was substantially shorter than that of the Amber ff99SBws-LJ trajectory, 1 *µ*s compared with 5 *µ*s, respectively. The Amber ff99SBws reconfiguration time shown in Fig. 5 should therefore be interpreted as a likely underestimate.

For the ProT*α*–protamine condensate at 128 mM KCl, no candidate lag passed the CK test for either force field. For Amber ff99SBws, the lowest CK score was 1.5 at a reference lag of 200 ns, and the local lag set used for averaging comprised 50, 100, 150, and 200 ns. For Amber ff99SBws-LJ, the lowest CK score was *S*_CK_ = 1.001 at a reference lag of 1000 ns. Although this value nominally failed the acceptance criterion, it exceeded the threshold by only 0.1%, indicating that the fitted model came very close to reproducing the longer-lag transition matrices within the bootstrap-estimated finite-sampling uncertainty. The result was nevertheless treated conservatively as a best-effort, not-converged estimate, and the local lag set used for averaging comprised 700, 800, and 1000 ns. The analyzed part of the Amber ff99SBws trajectory was substantially shorter than that of the Amber ff99SBws-LJ trajectory, 1.1 *µ*s compared with 7.8 *µ*s, respectively. The Amber ff99SBws reconfiguration time shown in Fig. 5 should therefore be interpreted as a likely underestimate.

For the ProT*α*–protamine condensate at 8 mM KCl, no candidate lag passed the CK test with either force field. For Amber ff99SBws, the lowest CK score was 1.5 at a reference lag of 300 ns, and the local lag set used for averaging comprised 200 and 300 ns. For Amber ff99SBws-LJ, the lowest CK score was 1.2 at a reference lag of 1500 ns, and the local lag set used for averaging comprised 1000 and 1500 ns. The analyzed part of the Amber ff99SBws trajectory was substantially shorter than that of the Amber ff99SBws-LJ trajectory, 1.3 *µ*s compared with 7.4 *µ*s, respectively. The Amber ff99SBws reconfiguration time shown in Fig. 5 should therefore be interpreted as a likely underestimate.

Importantly, irrespective of the exact choice of lag time, Amber ff99SBws-LJ agrees with the experimental reconfiguration times substantially better than Amber ff99SBws (Fig. S10).

## Data availability

The experimental osmometry data and optimized force-field parameter files obtained in this study are publicly available from Zenodo at https://zenodo.org/records/22101984.

## Acknowledgement

We thank Gerhard Hummer, Aritra Chowdhury and Rohan Eapen for insightful discussions, Annette Bieger Altermatt and Hille Ris for access to the osmometer and helpful discussions, Andrea Sottini for guidance on sample preparation and David Wang and George Pantelopulos for helpful comments on the manuscript. The work was supported by the Novo Nordisk Foundation (#NNF18OC0033926 to B.S.) and the Swiss National Science Foundation (grant number 10006187). We used the computational resources of Alps, Piz Daint and Eiger at the CSCS Swiss National Supercomputing Center, and of the National Institutes of Health HPC Biowulf cluster (http://hpc.nih.gov). R.B. was supported by the Intramural Research Program of the National Institute of Diabetes and Digestive and Kidney Diseases (NIDDK) within the National Institutes of Health (NIH). The contributions of the NIH author are considered works of the United States Government. The findings and conclusions presented in this paper are those of the author(s) and do not necessarily reflect the views of the NIH or the U.S. Department of Health and Human Services.

## Author contributions

M.T.I., R.B.B. and B.S. conceived the study. M.T.I. performed and analysed the simulations with the help of R.B.B. and input from B.S. M.T.I. performed the osmometry experiments and prepared the samples with assistance from V.v.R. and input from B.S. R.B.B. and B.S. supervised the project. M.T.I., R.B.B. and B.S. wrote the manuscript.

## References

(1) Holehouse, A. S.; Kragelund, B. B. The molecular basis for cellular function of intrinsically disordered protein regions. Nat. Rev. Mol. Cell Biol. 2024, 25, 187–211.

(2) Schuler, B.; Borgia, A.; Borgia, M. B.; Heidarsson, P. O.; Holmstrom, E. D.; Nettels, D.; Sottini, A. Binding without folding - the biomolecular function of disordered polyelectrolyte complexes. Curr. Opin. Struct. Biol. 2020, 60, 66–76.

(3) Borgia, A.; Borgia, M. B.; Bugge, K.; Kissling, V. M.; Heidarsson, P. O.; Fernandes, C. B.; Sottini, A.; Soranno, A.; Buholzer, K. J.; Nettels, D.; Kragelund, B. B.; Best, R. B.; Schuler, B. Extreme disorder in an ultrahigh-affinity protein complex. Nature 2018, 555, 61–66.

(4) Banani, S. F.; Lee, H. O.; Hyman, A. A.; Rosen, M. K. Biomolecular condensates: organizers of cellular biochemistry. Nat. Rev. Mol. Cell Biol. 2017, 18, 285–298.

(5) Brangwynne, C. P.; Eckmann, C. R.; Courson, D. S.; Rybarska, A.; Hoege, C.; Gharakhani, J.; Jülicher, F.; Hyman, A. A. Germline P granules are liquid droplets that localize by controlled dissolution/condensation. Science 2009, 324, 1729–1732.

(6) Cramer, P. Organization and regulation of gene transcription. Nature 2019, 573, 45–54.

(7) Bottaro, S.; Lindorff-Larsen, K. Biophysical experiments and biomolecular simulations: A perfect match? Science 2018, 361, 355–360.

(8) Wankowicz, S. A.; Bonomi, M. From possibility to precision in macromolecular ensemble prediction. Nat. Methods 2026, 1–9.

(9) Alberti, S. et al. Current practices in the study of biomolecular condensates: a community comment. Nat. Commun. 2025, 16, 7730.

(10) Dignon, G. L.; Zheng, W.; Kim, Y. C.; Best, R. B.; Mittal, J. Sequence determinants of protein phase behavior from a coarse-grained model. PLoS Comput. Biol. 2018, 14, e1005941.

(11) Alshareedah, I.; Borcherds, W. M.; Cohen, S. R.; Singh, A.; Posey, A. E.; Farag, M.; Bremer, A.; Strout, G. W.; Tomares, D. T.; Pappu, R. V.; Mittag, T.; Banerjee, P. R. Sequence-specific interactions determine viscoelasticity and ageing dynamics of protein condensates. Nat. Phys. 2024, 20, 1482–1491.

(12) Schäfer, L. V.; Stelzl, L. S. Deciphering driving forces of biomolecular phase separation from simulations. Curr. Opin. Struct. Biol. 2025, 92, 103026.

(13) Tesei, G.; Schulze, T. K.; Crehuet, R.; Lindorff-Larsen, K. Accurate model of liquid– liquid phase behavior of intrinsically disordered proteins from optimization of single-chain properties. Proc. Natl. Acad. Sci. U.S.A. 2021, 118, e2111696118.

(14) Gopi, S.; Qin, H.; Best, R. B.; Schuler, B. Beyond Equilibrium Ensembles: Time Rescaling in Coarse-Grained Simulations across Single-Molecule and Condensate Regimes. bioRxiv 2026, 2026–09.

(15) Galvanetto, N.; Ivanović, M. T.; Chowdhury, A.; Sottini, A.; Nüesch, M. F.; Nettels, D.; Best, R. B.; Schuler, B. Extreme dynamics in a biomolecular condensate. Nature 2023, 619, 876–883.

(16) Galvanetto, N.; Ivanović, M. T.; Del Grosso, S. A.; Chowdhury, A.; Sottini, A.; Nettels, D.; Best, R. B.; Schuler, B. Material properties of biomolecular condensates emerge from nanoscale dynamics. Proc. Natl. Acad. Sci. U.S.A. 2025, 122, e2424135122.

(17) Ivanović, M. T.; Best, R. B. All-atom simulations of biomolecular condensates. Curr. Opin. Struct. Biol. 2025, 93, 103101.

(18) Rauscher, S.; Gapsys, V.; Gajda, M. J.; Zweckstetter, M.; De Groot, B. L.; Grubmuller, H. Structural ensembles of intrinsically disordered proteins depend strongly on force field: a comparison to experiment. J. Chem. Theory Comput. 2015, 11, 5513– 5524.

(19) Best, R. B.; Hummer, G. Optimized molecular dynamics force fields applied to the helix- coil transition of polypeptides. J. Phys. Chem. B 2009, 113, 9004–9015.

(20) Best, R. B.; Zheng, W.; Mittal, J. Balanced protein–water interactions improve properties of disordered proteins and non-specific protein association. J. Chem. Theory Comput. 2014, 10, 5113–5124.

(21) Best, R. B.; Zhu, X.; Shim, J.; Lopes, P. E.; Mittal, J.; Feig, M.; MacKerell Jr, A. D. Optimization of the additive CHARMM all-atom protein force field targeting improved sampling of the backbone *ϕ*, *ψ* and side-chain *χ*1 and *χ*2 dihedral angles. J. Chem. Theory Comput. 2012, 8, 3257–3273.

(22) Shi, Y.; Xia, Z.; Zhang, J.; Best, R.; Wu, C.; Ponder, J. W.; Ren, P. Polarizable atomic multipole-based AMOEBA force field for proteins. J. Chem. Theory Comput. 2013, 9, 4046–4063.

(23) Piana, S.; Donchev, A. G.; Robustelli, P.; Shaw, D. E. Water dispersion interactions strongly influence simulated structural properties of disordered protein states. J. Phys. Chem. B 2015, 119, 5113–5123.

(24) Huang, J.; Rauscher, S.; Nawrocki, G.; Ran, T.; Feig, M.; De Groot, B. L.; Grubmüller, H.; MacKerell Jr, A. D. CHARMM36m: an improved force field for folded and intrinsically disordered proteins. Nat. Methods 2017, 14, 71–73.

(25) Robustelli, P.; Piana, S.; Shaw, D. E. Developing a molecular dynamics force field for both folded and disordered protein states. Proc. Natl. Acad. Sci. U.S.A. 2018, 115, E4758–E4766.

(26) Piana, S.; Robustelli, P.; Tan, D.; Chen, S.; Shaw, D. E. Development of a force field for the simulation of single-chain proteins and protein–protein complexes. J. Chem. Theory Comput. 2020, 16, 2494–2507.

(27) Lin, F.-Y.; Huang, J.; Pandey, P.; Rupakheti, C.; Li, J.; Roux, B.; MacKerell Jr, A. D. Further optimization and validation of the classical Drude polarizable protein force field. J. Chem. Theory Comput. 2020, 16, 3221–3239.

(28) Lindorff-Larsen, K.; Piana, S.; Dror, R. O.; Shaw, D. E. How fast-folding proteins fold. Science 2011, 334, 517–520.

(29) Henriques, J.; Cragnell, C.; Skepo, M. Molecular dynamics simulations of intrinsically disordered proteins: force field evaluation and comparison with experiment. J. Chem. Theory Comput. 2015, 11, 3420–3431.

(30) Shrestha, U. R.; Juneja, P.; Zhang, Q.; Gurumoorthy, V.; Borreguero, J. M.; Urban, V.; Cheng, X.; Pingali, S. V.; Smith, J. C.; O’Neill, H. M.; Petridis, L. Generation of the configurational ensemble of an intrinsically disordered protein from unbiased molecular dynamics simulation. Proc. Natl. Acad. Sci. U.S.A. 2019, 116, 20446–20452.

(31) Ahmed, M. C.; Papaleo, E.; Lindorff-Larsen, K. How well do force fields capture the strength of salt bridges in proteins? PeerJ 2018, 6, e4967.

(32) Mason, P. E.; Jungwirth, P.; Duboué-Dijon, E. Quantifying the strength of a salt bridge by neutron scattering and molecular dynamics. J. Phys. Chem. Lett. 2019, 10, 3254– 3259.

(33) Debiec, K. T.; Gronenborn, A. M.; Chong, L. T. Evaluating the strength of salt bridges: a comparison of current biomolecular force fields. J. Phys. Chem. B 2014, 118, 6561– 6569.

(34) Ivanović, M. T.; Holla, A.; Nüesch, M. F.; von Roten, V.; Schuler, B.; Best, R. B. Dynamical Buffering of Reconfiguration Dynamics in Intrinsically Disordered Proteins. JACS Au 2026, 6, 1900–1913.

(35) Lu, X.; Chen, J.; Huang, J. The continuous evolution of biomolecular force fields. Structure 2025, 33, 1138–1149.

(36) Leontyev, I.; Stuchebrukhov, A. Electronic continuum model for molecular dynamics simulations. J. Chem. Phys. 2009, 130, 085102.

(37) Kirby, B. J.; Jungwirth, P. Charge scaling manifesto: A way of reconciling the inherently macroscopic and microscopic natures of molecular simulations. J. Phys. Chem. Lett. 2019, 10, 7531–7536.

(38) Blazquez, S.; Conde, M.; Vega, C. Scaled charges for ions: An improvement but not the final word for modeling electrolytes in water. J. Chem. Phys. 2023, 158, 054505.

(39) Nencini, R.; Tempra, C.; Biriukov, D.; Riopedre-Fernandez, M.; Cruces Chamorro, V.; Polák, J.; Mason, P. E.; Ondo, D.; Heyda, J.; Ollila, O. H. S.; Jungwirth, P.; Javanainen, M.; Martinez-Seara, H. Effective inclusion of electronic polarization improves the description of electrostatic interactions: The prosECCo75 biomolecular force field. J. Chem. Theory Comput. 2024, 20, 7546–7559.

(40) Luo, Y.; Roux, B. Simulation of osmotic pressure in concentrated aqueous salt solutions. J. Phys. Chem. Lett. 2010, 1, 183–189.

(41) Miller, M. S.; Lay, W. K.; Elcock, A. H. Osmotic pressure simulations of amino acids and peptides highlight potential routes to protein force field parameterization. J. Phys. Chem. B 2016, 120, 8217–8229.

(42) Miller, M. S.; Lay, W. K.; Li, S.; Hacker, W. C.; An, J.; Ren, J.; Elcock, A. H. Reparametrization of protein force field nonbonded interactions guided by osmotic coefficient measurements from molecular dynamics simulations. J. Chem. Theory Comput. 2017, 13, 1812–1826.

(43) Hagler, A.; Huler, E.; Lifson, S. Energy functions for peptides and proteins. I. Derivation of a consistent force field including the hydrogen bond from amide crystals. J. Am. Chem. Soc. 1974, 96, 5319–5327.

(44) Bonner, O. D. Osmotic and activity coefficients of sodium and potassium glutamate at 298.15 K. J. Chem. Eng. Data 1981, 26, 147–148.

(45) Bonner, O. D. Osmotic and activity coefficients of some amino acids and their hydrochloride salts at 298.15 K. J. Chem. Eng. Data 1982, 27, 422–423.

(46) Robinson, R. A.; Stokes, R. H. Electrolyte solutions; Courier Corporation, 2002.

(47) Fisher, R. S.; Elbaum-Garfinkle, S. Tunable multiphase dynamics of arginine and lysine liquid condensates. Nat. Commun. 2020, 11, 4628.

(48) Schuster, B. S.; Dignon, G. L.; Tang, W. S.; Kelley, F. M.; Ranganath, A. K.; Jahnke, C. N.; Simpkins, A. G.; Regy, R. M.; Hammer, D. A.; Good, M. C.; Mittal, J. Identifying sequence perturbations to an intrinsically disordered protein that determine its phase-separation behavior. Proc. Natl. Acad. Sci. U.S.A. 2020, 117, 11421–11431.

(49) von Roten, V.; Ivanović, M. T.; Gopi, S.; Holla, A.; Prestel, A.; Nüesch, M.; Tamburrini, K. C.; Nettels, D.; Kragelund, B. B.; Best, R. B.; Schuler, B. Cooperativity, dynamics, and the free-energy surfaces of charge-patterned IDPs. bioRxiv 2026, 2026– 05.

(50) Abascal, J. L.; Vega, C. A general purpose model for the condensed phases of water: TIP4P/2005. J. Chem. Phys. 2005, 123, 234505.

(51) Smith, P. K.; Smith, E. R. Thermodynamic properties of solutions of amino acids and related substances: II. The activity of aliphatic amino acids in aqueous solution at twenty-five degrees. J. Biol. Chem. 1937, 121, 607–613.

(52) Smith, E. R.; Smith, P. K. Thermodynamic Properties of Solutions of Amino Acids and Related Substances: VI. The Activities of Some Peptides in Aqueous Solution at Twenty-Five Degrees. J. Biol. Chem. 1940, 135, 273–279.

(53) Kann, Z.; Skinner, J. A scaled-ionic-charge simulation model that reproduces enhanced and suppressed water diffusion in aqueous salt solutions. J. Chem. Phys. 2014, 141, 104507.

(54) Fuentes-Azcatl, R.; Barbosa, M. C. Sodium chloride, NaCl/*ɛ*: New force field. J. Phys. Chem. B 2016, 120, 2460–2470.

(55) Duboué-Dijon, E.; Mason, P. E.; Fischer, H. E.; Jungwirth, P. Hydration and ion pairing in aqueous Mg2+ and Zn2+ solutions: force-field description aided by neutron scattering experiments and ab initio molecular dynamics simulations. J. Phys. Chem. B 2017, 122, 3296–3306.

(56) Duboue-Dijon, E.; Javanainen, M.; Delcroix, P.; Jungwirth, P.; Martinez-Seara, H. A practical guide to biologically relevant molecular simulations with charge scaling for electronic polarization. J. Chem. Phys. 2020, 153, 050901.

(57) Ohtaki, H.; Fukushima, N. A structural study of saturated aqueous solutions of some alkali halides by X-ray diffraction. J. Solut. Chem. 1992, 21, 23–38.

(58) Marcus, Y. Thermodynamics of solvation of ions. Part 5.—Gibbs free energy of hydration at 298.15 K. J. Chem. Soc., Faraday Trans. 1991, 87, 2995–2999.

(59) Holla, A.; Martin, E. W.; Dannenhoffer-Lafage, T.; Ruff, K. M.; König, S. L. B.; Nüesch, M. F.; Chowdhury, A.; Louis, J. M.; Soranno, A.; Nettels, D.; Pappu, R. V.; Best, R. B.; Mittag, T.; Schuler, B. Identifying sequence effects on chain dimensions of disordered proteins by integrating experiments and simulations. JACS Au 2024, 4, 4729–4743.

(60) Bugge, K.; Sottini, A.; Ivanović, M. T.; Buus, F. S.; Saar, D.; Fernandes, C. B.; Kocher, F.; Martinsen, J. H.; Schuler, B.; Best, R. B.; Kragelund, B. B. Role of charges in a dynamic disordered complex between an IDP and a folded domain. Nat. Commun. 2025, 16, 3242.

(61) Martinsen, J. H.; Saar, D.; Fernandes, C. B.; Schuler, B.; Bugge, K.; Kragelund, B. B. Structure, dynamics, and stability of the globular domain of human linker histone H1.0 and the role of positive charges. Protein Sci. 2022, 31, 918–932.

(62) Sabari, B. R. et al. Coactivator condensation at super-enhancers links phase separation and gene control. Science 2018, 361, eaar3958.

(63) Shin, Y.; Chang, Y.-C.; Lee, D. S.; Berry, J.; Sanders, D. W.; Ronceray, P.; Wingreen, N. S.; Haataja, M.; Brangwynne, C. P. Liquid nuclear condensates mechanically sense and restructure the genome. Cell 2018, 175, 1481–1491.

(64) Yoo, J.; Aksimentiev, A. Improved parameterization of amine–carboxylate and amine– phosphate interactions for molecular dynamics simulations using the CHARMM and AMBER force fields. J. Chem. Theory Comput. 2016, 12, 430–443.

(65) Cornell, W. D.; Cieplak, P.; Bayly, C. I.; Gould, I. R.; Merz, K. M.; Ferguson, D. M.; Spellmeyer, D. C.; Fox, T.; Caldwell, J. W.; Kollman, P. A. A second generation force field for the simulation of proteins, nucleic acids, and organic molecules. J. Am. Chem. Soc. 1995, 117, 5179–5197.

(66) Best, R. B.; de Sancho, D.; Mittal, J. Residue-specific *α*-helix propensities from molecular simulation. Biophys. J. 2012, 102, 1462–1467.

(67) Mahmoudi, S.; Gruene, T.; Schröder, C.; Ferjaoui, K. D.; Fröjdh, E.; Mozzanica, A.; Takaba, K.; Volkov, A.; Maisriml, J.; Paunović, V.; van Bokhoven, J. A.; Keppler, B. K. Experimental determination of partial charges with electron diffraction. Nature 2025, 645, 88–94.

(68) Bayly, C. I.; Cieplak, P.; Cornell, W.; Kollman, P. A. A well-behaved electrostatic potential based method using charge restraints for deriving atomic charges: the RESP model. J. Phys. Chem. 1993, 97, 10269–10280.

(69) Gaussian, Inc. Gaussian 16, Revision C.01. 2016.

(70) Abraham, M. J.; Murtola, T.; Schulz, R.; Páll, S.; Smith, J. C.; Hess, B.; Lindahl, E. GROMACS: High performance molecular simulations through multi-level parallelism from laptops to supercomputers. SoftwareX 2015, 1, 19–25.

(71) Parrinello, M.; Rahman, A. Polymorphic transitions in single crystals: A new molecular dynamics method. J. Appl. Phys. 1981, 52, 7182–7190.

(72) Bussi, G.; Donadio, D.; Parrinello, M. Canonical sampling through velocity rescaling. J. Chem. Phys. 2007, 126, 014101.

(73) Darden, T.; York, D.; Pedersen, L. Particle mesh Ewald: An N log (N) method for Ewald sums in large systems. J. Chem. Phys. 1993, 98, 10089–10092.

(74) Hess, B.; Bekker, H.; Berendsen, H. J.; Fraaije, J. G. LINCS: A linear constraint solver for molecular simulations. J. Comput. Chem. 1997, 18, 1463–1472.

(75) Satarifard, V.; Kashefolgheta, S.; Vila Verde, A.; Grafmüller, A. Is the solution activity derivative sufficient to parametrize ion–ion interactions? Ions for tip5p water. J. Chem. Theory Comput. 2017, 13, 2112–2122.

(76) Klose, D.; Holla, A.; Gmeiner, C.; Nettels, D.; Ritsch, I.; Bross, N.; Yulikov, M.; Allain, F. H.-T.; Schuler, B.; Jeschke, G. Resolving distance variations by single-molecule FRET and EPR spectroscopy using rotamer libraries. Biophys. J. 2021, 120, 4842– 4858.

(77) Best, R. B.; Hofmann, H.; Nettels, D.; Schuler, B. Quantitative interpretation of FRET experiments via molecular simulation: force field and validation. Biophys. J. 2015, 108, 2721–2731.

(78) Nüesch, M.; Ivanović, M. T.; Nettels, D.; Best, R. B.; Schuler, B. Accuracy of distance distributions and dynamics from single-molecule FRET. Biophys. J. 2025, 124, 3408– 3427.

(79) Best, R. B.; Hummer, G.; Eaton, W. A. Native contacts determine protein folding mechanisms in atomistic simulations. Proc. Natl. Acad. Sci. U.S.A. 2013, 110, 17874–17879.

(80) Hummer, G. Position-dependent diffusion coefficients and free energies from Bayesian analysis of equilibrium and replica molecular dynamics simulations. New J. Phys. 2005, 7, 34–34.

(81) Bicout, D.; Szabo, A. Electron transfer reaction dynamics in non-Debye solvents. J. Chem. Phys. 1998, 109, 2325–2338.

(82) Prinz, J.-H.; Wu, H.; Sarich, M.; Keller, B.; Senne, M.; Held, M.; Chodera, J. D.; Schütte, C.; Noé, F. Markov models of molecular kinetics: Generation and validation. J. Chem. Phys. 2011, 134, 174105.

