## Supplementary Information for "Predictive all-atom simulations of disordered proteins and biomolecular condensates through osmometry-guided force-field optimization"

### Supporting Figures

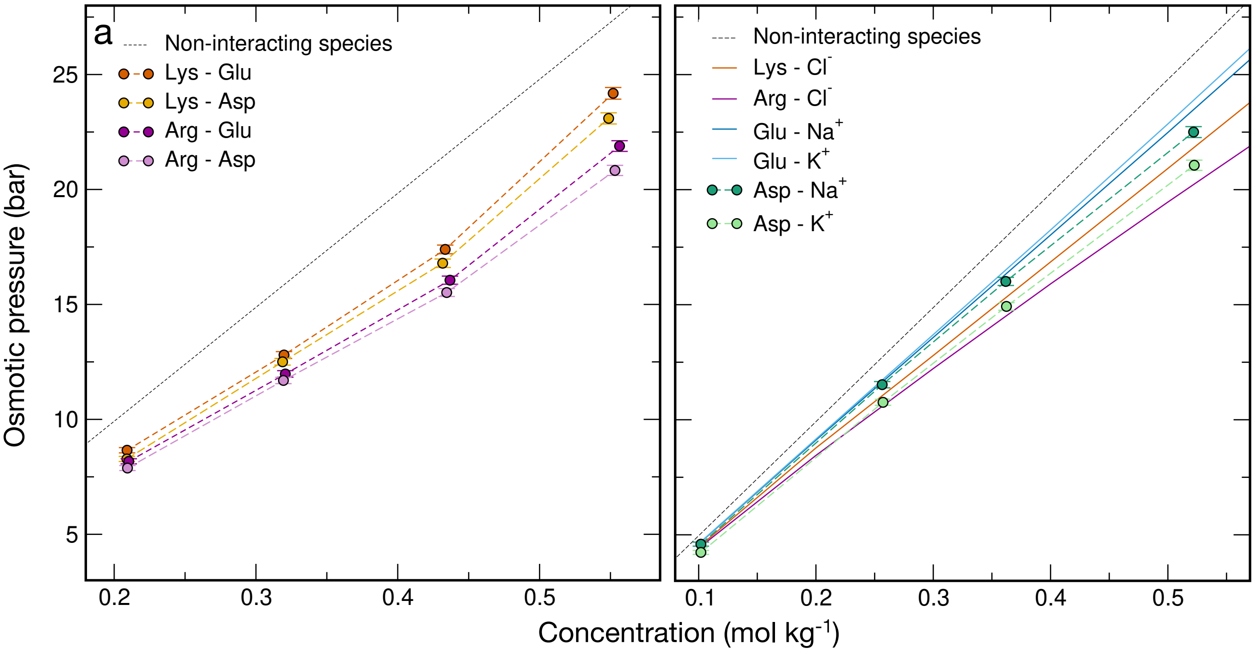


**Figure S1: Experimental osmotic pressure values.** (a) Residue–residue interactions; (b) residue–ion interactions. Theoretical osmotic pressures for pairs of non-interacting species are shown versus molal concentration as black dotted lines. All measured osmotic pressures lie below the corresponding non-interacting values, consistent with predominantly attractive interactions in these solutions of oppositely charged species. Symbols and dashed lines represent data from this work and solid lines represent data taken from the literature.[1–3]

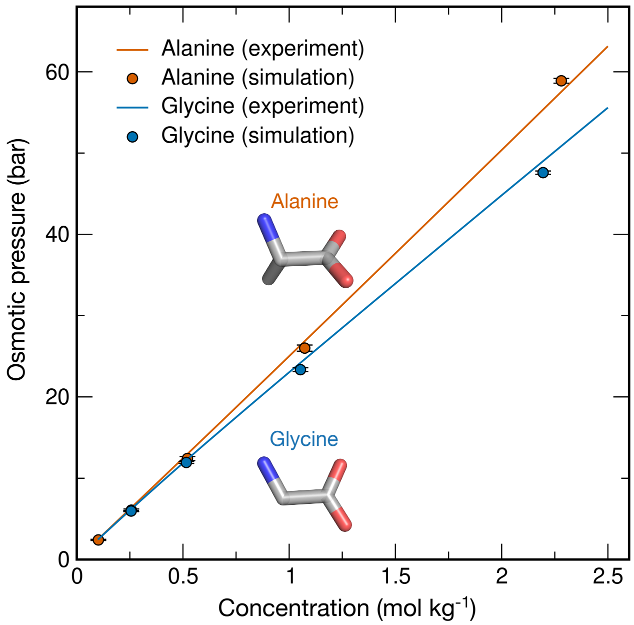


**Figure S2: Zwitterionic parameter validation.** Experimental osmotic pressures (solid lines) and osmotic pressures calculated from simulations of alanine and glycine in their zwitterionic forms (filled circles).

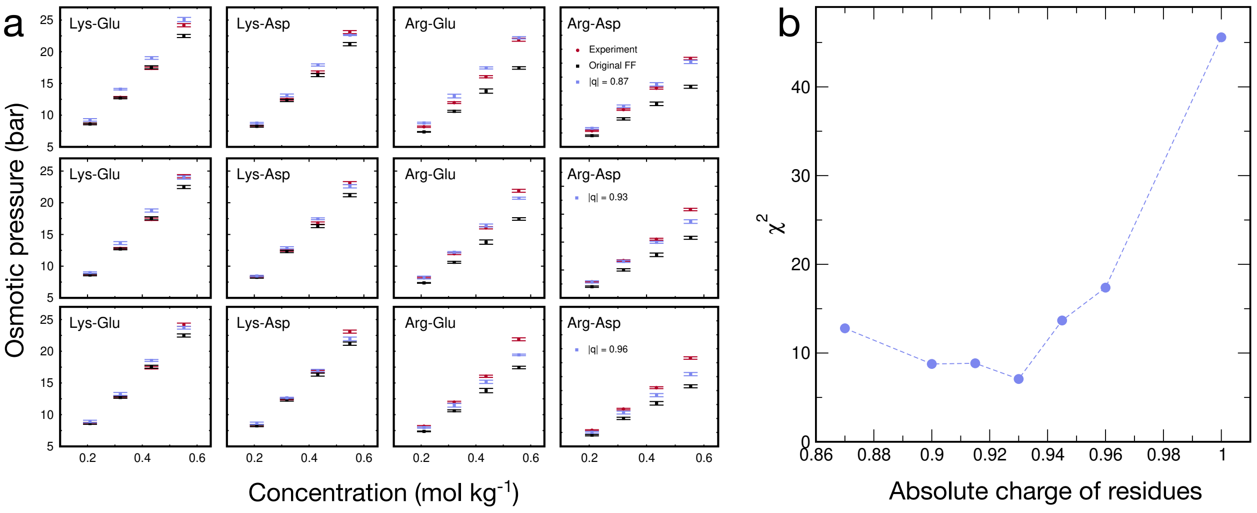


**Figure S3: Amber ff99SBws-CS: Determining absolute charge of residues.** We assumed that the absolute charge of all charged residues is scaled to the same value. (a) Osmotic pressure of four sets of simulations (columns: Lys-Glu, Lys-Asp, Arg-Glu and Arg-Asp) each at four different amino acid concentrations, and their comparison with experiment. Examples are shown for the absolute charges scaled to ± 0.87 (top row), ± 0.93 (middle row) and ± 0.96 (bottom row). (b) Overall agreement with experimental osmotic pressure data as a function of selected absolute charge, expressed using the unreduced $\chi^2$ parameter.

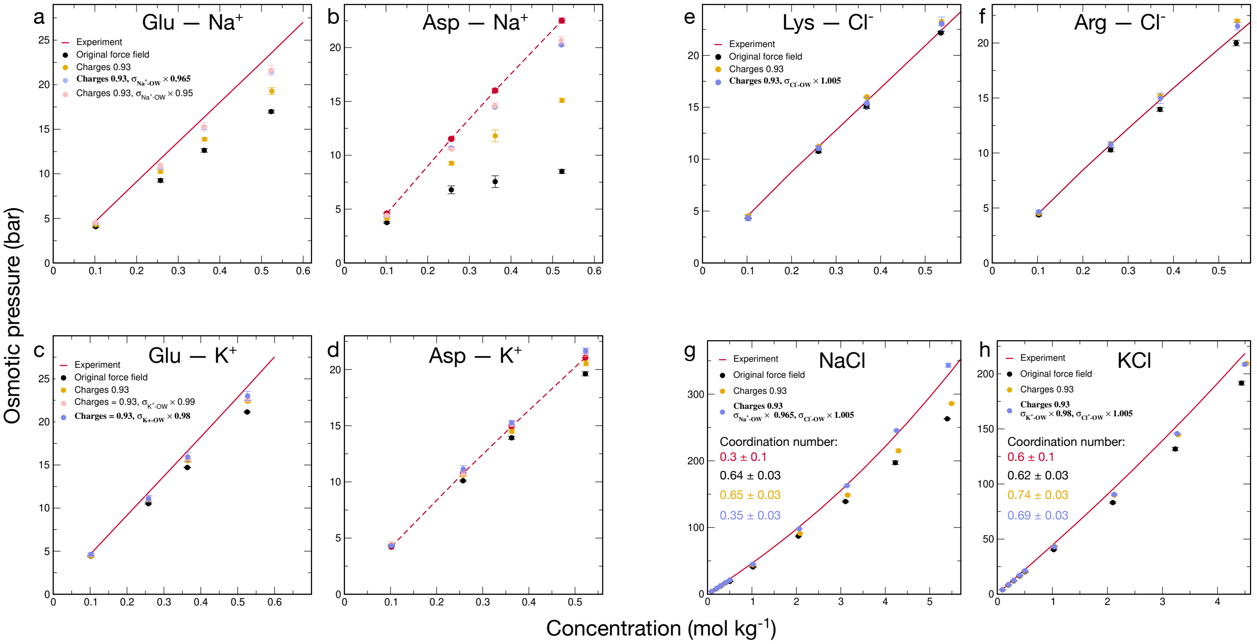


**Figure S4: Amber ff99SBws-CS optimization: Optimizing protein–ion and ion–ion interactions after scaling the charges.** (a,b) Glu–$Na^+$ and Asp–$Na^+$ interactions are too strong with the original force field, as reported previously for several force fields.[4–6] Scaling the charges of the charged residues and $Na^+$ to ±0.93 increases the osmotic pressure toward the experimental values, but the interactions remain too strong. Further improvement is obtained by modifying the Lennard–Jones $\sigma$ parameter between $Na^+$ and water oxygen (OW), as indicated in the panel legends. (c,d) The corresponding optimization is shown for Glu–$K^+$ and Asp–$K^+$, where adjustment of the $K^+$–OW Lennard–Jones $\sigma$ parameter improves agreement with experiment. (e,f) Lys–$Cl^-$ and Arg–$Cl^-$ interactions are already described well after charge scaling; a small adjustment of the $Cl^-$–OW Lennard–Jones $\sigma$ parameter retains this agreement. (g,h) NaCl and KCl osmotic pressures are shown for the same parameter sets used for the residue–ion optimization. $K^+$–$Cl^-$ interactions are slightly too strong with the original force field.[6] For the experimental data, solid lines denote values taken from the literature,[1,2] whereas dashed lines denote measurements obtained in this work. The corresponding experimental and simulated coordination numbers are reported within the panels. The final parameter set is highlighted in bold in each panel.

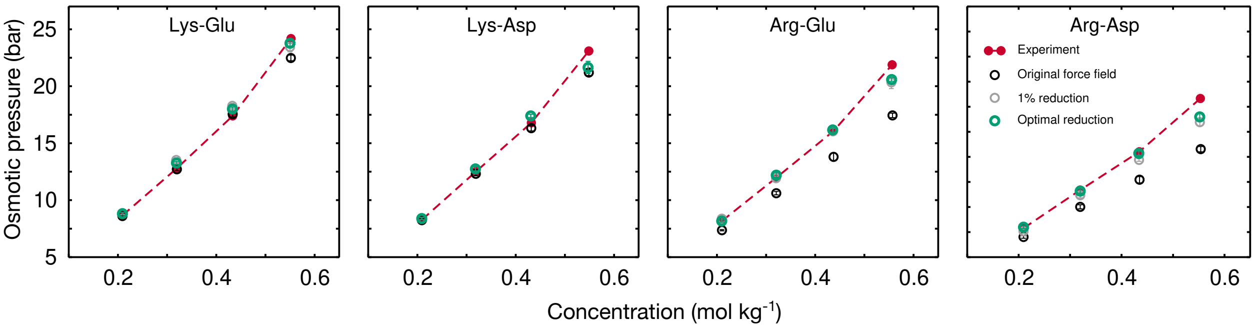


**Figure S5: Amber ff99SBws-LJ optimization: effect of reducing the side-chain–OW Lennard-Jones $\sigma$ parameters on the osmotic pressures of residue–residue mixtures.** Experimental osmotic pressures (red circles and dashed lines) are compared with simulations using Amber ff99SBws (black circles), a uniform 1% reduction of the side-chain–OW $\sigma$ parameters for all four charged residues (gray circles), and optimized residue-specific reductions (green circles). The corresponding optimized scaling factors are 0.99 for $\sigma_{\mathrm{Lys-OW}}$, 0.985 for $\sigma_{\mathrm{Arg-OW}}$, 0.995 for $\sigma_{\mathrm{Glu-OW}}$, and 0.99 for $\sigma_{\mathrm{Asp-OW}}$.

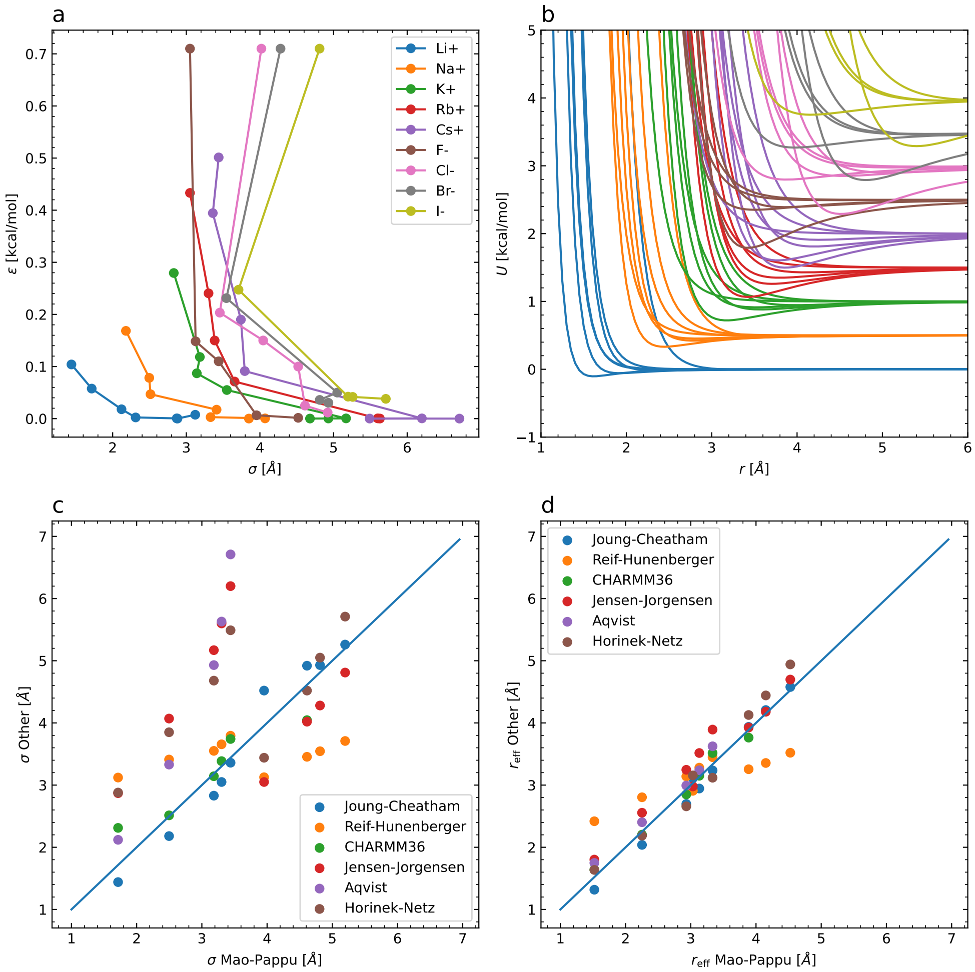


**Figure S6: Effective ion radius in additive force fields** (a) Anti-correlation between Lennard-Jones $\sigma$ and $\epsilon$ for monovalent ion parameters from Mao and Pappu,[7] Joung and Cheatham,[8] Reif and Hünenberger ($M_E$ parameters),[9] CHARMM36,[10] Jensen and Jorgensen,[11] Aqvist[12] and Horinek et al.[13] (b) Corresponding Lennard-Jones potentials (colors as in (a)). (c) Correlation between Mao-Pappu parameters for $\sigma$ and those from other force fields (see Legend). (d) Correlation between Mao-Pappu effective radii $r_{\text{eff}}$ (as defined in main text) and those from other force fields.

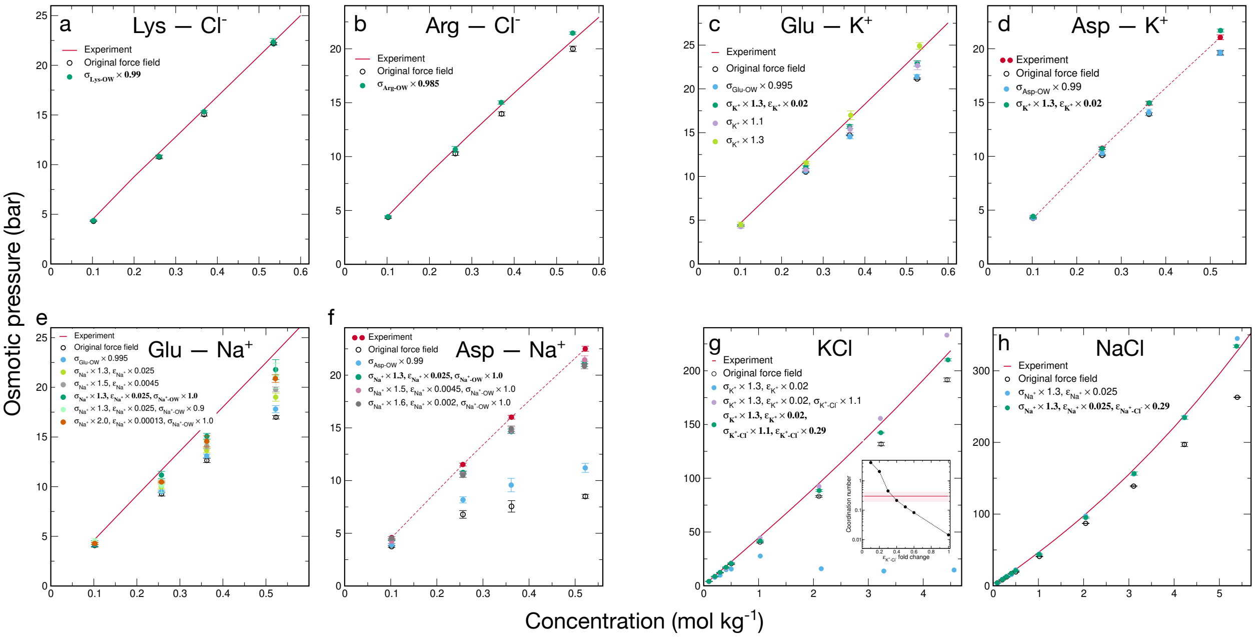

**Figure S7: Amber ff99SBws-LJ: protein–ion and ion–ion parameters.** (a,b) Lys/Arg–$Cl^-$ interactions: experimental osmotic pressure (red), osmotic pressure calculated from simulations using the original force field (black), and osmotic pressure after Lys/Arg–OW interactions have been modified (green). (c,d) Glu/Asp–$K^+$ interactions: experimental osmotic pressure (red), osmotic pressure calculated from simulations using the original force field (black), and osmotic pressure after Glu/Asp–OW interactions have been modified (light blue). Because Glu/Asp–$K^+$ interactions remained too strong after Glu/Asp–OW modifications, we further adjusted the $K^+$ Lennard-Jones $\sigma$ and $\epsilon$ such that the effective radius of $K^+$ is changed by less than 2%. The final modification, with $K^+$ $\sigma$ increased by a factor of 1.3 and $K^+$ $\epsilon$ scaled by a factor of 0.02 (dark green), provides a compromise that on average describes both Glu–$K^+$ and Asp–$K^+$ interactions well. The effect of changing $\sigma$ can be seen by comparing the light blue, purple, and light green symbols in panel (c), whereas the effect of changing $\epsilon$ can be seen by comparing the light green and dark green symbols in panel (c). Note that for the dark green, light green, and purple symbols, Glu–OW $\sigma$ was also modified by a factor of 0.995, as explicitly indicated only in the light blue symbol legend. (e,f) Glu/Asp–$Na^+$ interactions: experimental osmotic pressure (red), osmotic pressure calculated from simulations using the original force field (black), and osmotic pressure after Glu/Asp–OW interactions have been modified (light blue). Note that for all symbols listed below the light blue entry in the legend, Glu/Asp–OW interactions have been modified as described in the main text and in Fig. S5 (reducing Glu–OW $\sigma$ by 0.5% and Asp–OW $\sigma$ by 1%). As apparent from panels (e) and (f), increasing the $Na^+$ $\sigma$ improves agreement with experiment, but this improvement nearly saturates when $\sigma$ has been increased by a factor of 1.3. The maximal improvement is obtained for a $Na^+$–OW $\sigma$ value corresponding to the $Na^+$–OW $\sigma$ of the unmodified ion (i.e., the value before $Na^+$ $\sigma$ and $\epsilon$ were changed, denoted by Na-OW $\sigma$ 1.0). Further modifications of $Na^+$–OW $\sigma$ do not lead to any noticeable improvement. (g) $K^+$–$Cl^-$ interactions: experimental osmotic pressure (red) and osmotic pressure calculated from simulations using the original force field (black). The osmotic pressure obtained from simulations in which the $K^+$ Lennard-Jones parameters were changed as in the Glu/Asp–$K^+$ optimization is shown in light blue. This change makes $K^+$–$Cl^-$ interactions too strong, which is reflected in an osmotic pressure that is too low. To correct this, we modified the $K^+$–$Cl^-$ Lennard-Jones parameters. Increasing $K^+$–$Cl^-$ $\sigma$ weakens $K^+$–$Cl^-$ interactions, as seen from the increase in osmotic pressure (purple symbols). After optimizing $K^+$–$Cl^-$ $\sigma$, we also adjusted $K^+$–$Cl^-$ $\epsilon$ to match the experimental $K^+$–$Cl^-$ coordination number at near-saturation concentration (inset). Decreasing $K^+$–$Cl^-$ $\epsilon$ moderately decreases the calculated osmotic pressure (green symbols) while bringing the coordination number into the experimental range. (h) $Na^+$–$Cl^-$ interactions: experimental osmotic pressure (red) and osmotic pressure calculated from simulations using the original force field (black). The osmotic pressure obtained from simulations in which the $Na^+$ Lennard-Jones parameters were changed as in the Glu/Asp–$Na^+$ optimization is shown in light blue. In contrast to $K^+$–$Cl^-$, where increasing the $K^+$ size makes $K^+$–$Cl^-$ interactions too favorable, increasing the $Na^+$ size improves agreement with the experimental osmotic pressure. To obtain a $Na^+$–$Cl^-$ coordination number at near-saturation concentration that agrees with experiment, we further modified $Na^+$–$Cl^-$ $\epsilon$ (green symbols). For the experimental data, solid lines denote values taken from the literature,[1,2] whereas dashed lines denote measurements obtained in this work. The final simulation parameter set is highlighted in bold in each panel.

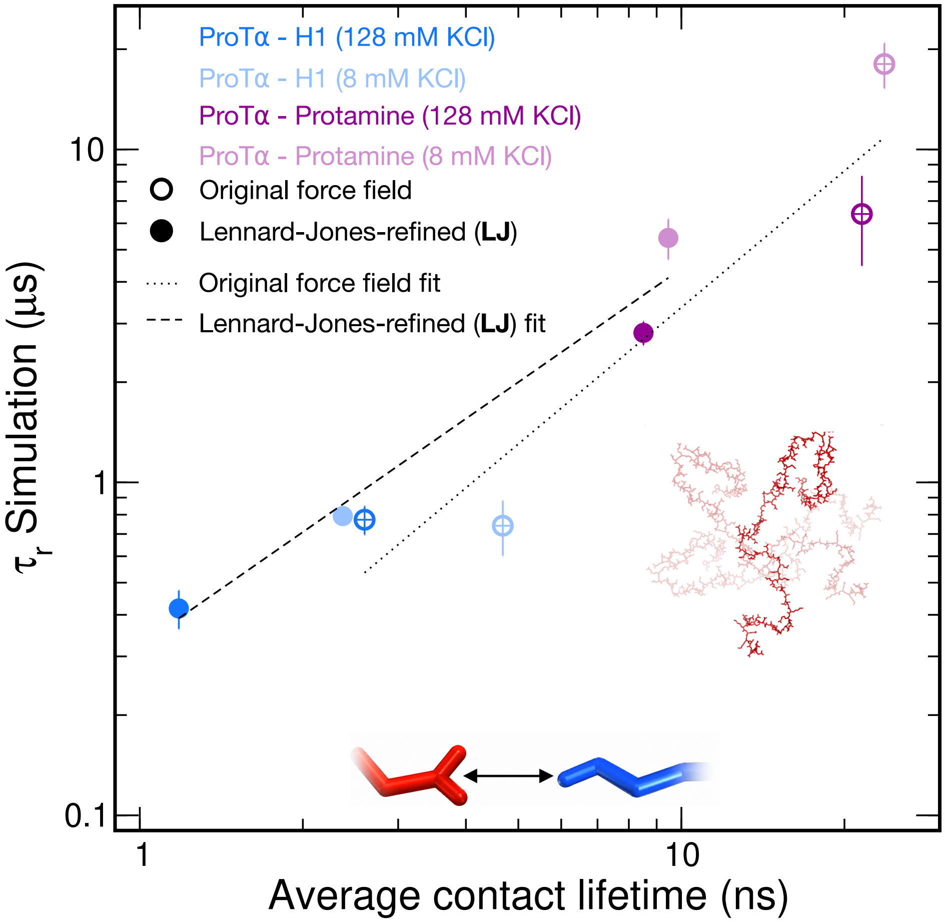


**Figure S8: Relation between intermolecular residue–residue contact lifetimes and ProT$\alpha$ chain reconfiguration times in condensates.** ProT$\alpha$ reconfiguration times calculated from the simulations are shown as a function of the mean lifetime of intermolecular residue–residue contacts involving ProT$\alpha$. Contact lifetimes were averaged over all ProT$\alpha$ residues and all 96 ProT$\alpha$ chains in each simulation. Dotted and dashed lines show separate log–log fits to the Amber ff99SBws and Amber ff99SBws-LJ data, respectively. For each force field, the four condensate data points were fitted by ordinary least squares after transforming both variables to $\log_{10}$ scale, that is, $\log_{10} y = a \log_{10} x + b$. Contact lifetimes and chain reconfiguration times are strongly correlated across the four condensates for both force fields. For every condensate, Amber ff99SBws-LJ gives shorter contact lifetimes and shorter reconfiguration times than Amber ff99SBws, with the reconfiguration times in substantially better agreement with experiment (Fig. 5).

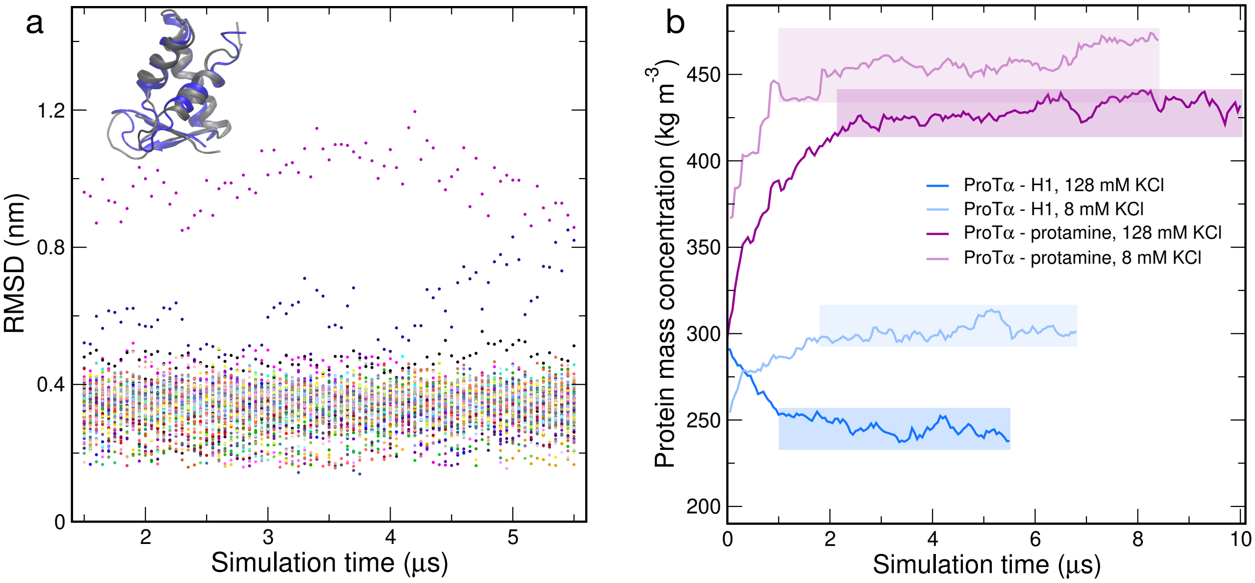


**Figure S9: Globular domain stability and protein mass concentration in the condensates as a function of simulation time.** (a) Stability of the H1 globular domains (GDs) in the ProT$\alpha$–H1 condensate at 128 mM KCl with Amber ff99SBws-LJ, quantified as the backbone RMSD relative to the experimental structure (PDB 6HQ1).[14] RMSDs for each of the 80 GDs were calculated every 50 ns. Only 4 of 80 GDs (5%) had a time-averaged RMSD above 0.4 nm, in line with the fraction of partially unfolded domains observed previously with Amber ff99SBws and with the experimentally determined stability in dilute solution.[15] Backbone RMSDs of 0.2–0.4 nm can largely be attributed to the flexibility of loops in the domain, illustrated by the superposition of the experimental structure and a simulated GD structure with an RMSD of 0.4 nm (inset). (b) Protein mass concentrations in the dense phases were calculated in 50 ns blocks. Initial trajectory segments during which the protein concentration changed substantially were treated as equilibration and excluded from all analyses. Shaded regions indicate the trajectory segments used for the analyses reported in the manuscript. Amber ff99SBws-LJ gives lower mean dense-phase protein concentrations than Amber ff99SBws in all four systems. The concentrations decreased from $(290 \pm 10)\,\mathrm{kg\,m^{-3}}$ to $(245 \pm 5)\,\mathrm{kg\,m^{-3}}$ for ProT$\alpha$–H1 at 128 mM KCl,[15] from $(339 \pm 5)\,\mathrm{kg\,m^{-3}}$ to $(301 \pm 5)\,\mathrm{kg\,m^{-3}}$ for ProT$\alpha$–H1 at 8 mM KCl,[16] from $(460 \pm 3)\,\mathrm{kg\,m^{-3}}$ to $(429 \pm 6)\,\mathrm{kg\,m^{-3}}$ for ProT$\alpha$–protamine at 128 mM KCl,[16] and from $(473 \pm 6)\,\mathrm{kg\,m^{-3}}$ to $(455 \pm 4)\,\mathrm{kg\,m^{-3}}$ for ProT$\alpha$–protamine at 8 mM KCl.[16] In each comparison, the first value was obtained with Amber ff99SBws and the second with Amber ff99SBws-LJ. These changes correspond to decreases of approximately 16%, 11%, 7%, and 4%, respectively. For the protamine-containing condensates, the dense-phase protein concentration differs by only 4–7% between the two parameter sets, whereas the chain reconfiguration times differ by approximately threefold (Fig. 5 and Table S2), suggesting that dense-phase protein concentration is not the only factor determining IDP conformational dynamics within condensates.

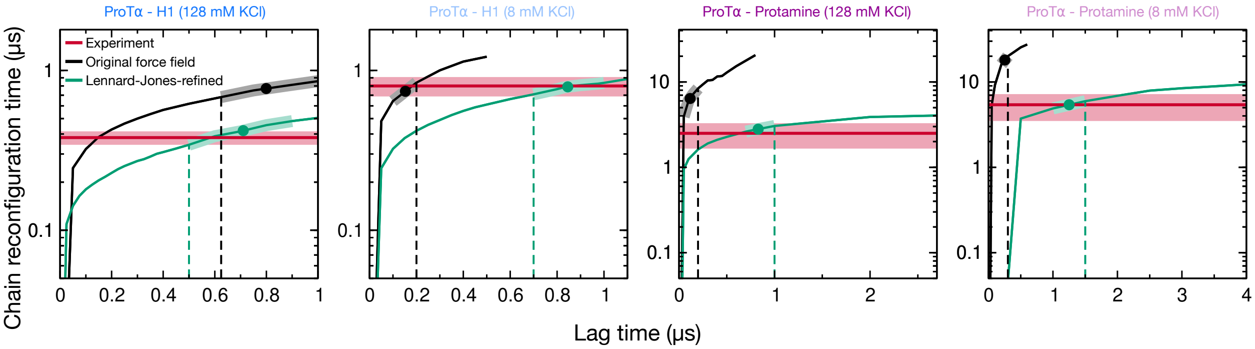


**Figure S10: ProT$\alpha$ chain reconfiguration time as a function of diffusion model lag time.** For each simulated condensate, the reconfiguration time of ProT$\alpha$ is shown as a function of the lag time used to construct a one-dimensional diffusion model from the distances between residues 58 and 112 of all ProT$\alpha$ chains. Black and green curves show results obtained with Amber ff99SBws and Amber ff99SBws-LJ, respectively. Vertical dashed lines mark the selected lag for each simulation: the smallest candidate lag satisfying the Chapman–Kolmogorov (CK) criterion or, when no candidate passed, the lag with the lowest CK score. Shaded curve segments indicate the lag-time windows used for averaging. The reported simulated reconfiguration time is the unweighted mean of the reconfiguration times calculated at the candidate lags within each window, and its uncertainty is their sample standard deviation. Filled circles show these mean values. Because an averaged value does not itself have an associated lag, each circle is positioned horizontally by interpolation along the corresponding curve; this position is used only for display. Experimental reconfiguration times and their uncertainties are shown by the red lines and shaded bands, respectively. Further details of the lag-time selection and CK analysis are provided in Methods.

### Supporting Tables

Table S1: **Protein and IDR sequences used in simulations.** The 16 linker IDRs are 68 residues long and contain Cys residues at positions 5 and 63 for site-specific dye labelling. G denotes Cy3B (green donor; net charge $Q = 0$) and R denotes CF660R (red acceptor; net charge $Q = -1$). GR indicates Cy3B at position 5 and CF660R at position 63, whereas RG indicates the reverse labelling orientation. The dyes were represented explicitly in the linker-IDR simulations. A dash indicates simulations without explicitly represented fluorophores.

| Protein/IDR | Sequence | Labelling |
|---|---|---|
| *Linker IDRs* | | |
| sGrich | GSGSCSGGYGSERGGGYGSERGGGYGSERGGGYGSERGGGYGSQRSGGGYGGSQRSSYGSGSCTLGPR | GR |
| d$\kappa$h | GSGSCAMGGGPGPGTDFTSDQADFPDTLFQEFEPPAPRPGLAGGRPGDAALLSAAYGRRRLLCTLGPR | GR |
| dGrich | GSGSCGPRTGLEGAGMAGGSGQQKRVFDGQSGPQDLGEAYRPLNHDGGDGGNRYSVIDRIQECTLGPR | GR |
| sPTBP | GSGSCRPDLPSGDSQPSLDQTMAAAFGLSVPNVHGALAPLAIPSAAAAAAAAGRIAIPGLAGCTLGPR | RG |
| sCh | GSGSCKKAQPEMNDKDDNESGNEDAEENHDDEEDENEEEDRQVDQASKNKESKRKAQNKREDCTLGPR | GR |
| sNrich | GSGSCLDQEDNNGPLLIKTANNLIQNNSNMLPLNALHNAPPMHLNEGGISNMRVNDSLPSNTCTLGPR | GR |
| s$\kappa$h | GSGSCKPRRLSKLRRSKKPADEENNAASQDPTVEATQERGQASEDPENAANNAKQAKPTSDDCTLGPR | RG |
| dCh+ | GSGSCQTPLKRIKVKTPGKSGAAAREGSVVSGTDGPTQTGKPERRKRLNPPKDKLIDMDDADCTLGPR | RG |
| dCh- | GSGSCMGLPTGMEEKEEGTDESEQKPVVQTPAQPDDSAEVDSAALDQAESAKQQGPILTKHGCTLGPR | RG |
| s$\kappa$l | GSGSCAPEGVFKLPAPPKEIKKSEKSEKSSDSINDKKEVESTTKTAATTTTTKKGTDNNTQFCTLGPR | RG |
| sNh+ | GSGSCPEEIETRKKDRKNRREMLKRTSALQPAAPKPTHKKPVPKRNVGAERKSTINEDLLPPCTLGPR | GR |
| dArich | GSGSCLPTGAEGRDSSKGEDSAEETEAKPAVVAPAPVVEAVSTPSAAFPSDATAEQGPILTKCTLGPR | RG |
| dErich | GSGSCEGEMFGVMSEENSVVLSVEQPAELKEVADVSPPPTTRNHTIEMKPPLSAQQSESNPVCTLGPR | RG |
| dTRBP | GSGSCLEPALEDSSSFSPLDSSLPEDIPVFTAAAAATPVPSVVLTRSPPMELQPPVSPQQSECTLGPR | RG |
| d$\kappa$l | GSGSCLPTGMDQKTTELMKVDEPVSTQETLPPVIKMEPEPVPITETPTDENARQQGPILTKHCTLGPR | GR |
| sNh- | GSGSCKKKLQDTPSEPMEKDPAEPETVPDGETPEDENPTEEGADNSSAKMEEEEEEEEEEEEEECTLGPR | RG |
| *Proteins used in condensate simulations* | | |
| ProT$\alpha$ | GPMSDAAVDTSSEITTKDLKEKKEVVEEAENGRDAPANGNANEENGEQEADNEVDEEEEEGGEEEEEEEEGDGEEEDGDE DEEAESATGKRAAEDDEDDDVDTKKQKTDEDD | – |
| H1 | TENSTSAPAAKPKRAKASKKSTDHPKYSDMIVAAIQAEKNRAGSSRQSIQKYIKSHYKVGENADSQIKLSIKRLVTTGVL KQTKGVGASGSFRLAKSDEPKKSVAFKKTKKEIKKVATPKKASKPKKAASKAPTKKPKATPVKKAKKKLAATPKKAKKPK TVKAKPVKASKPKKAKPVKPKAKSSAKRAGKKK | – |
| Protamine | MPRRRRSSSRPVRRRRRPRVSRRRRRRGGRRRR | – |

Table S2: Mean FRET efficiencies, $\langle E \rangle$, and ProT$\alpha$ chain reconfiguration times, $\tau_r$, are shown for ProT$\alpha$–H1 and ProT$\alpha$–protamine condensates at the 8 and 128 mM KCl simulation conditions. Experiments were performed in TEK buffer, which contributes approximately 8 mM to the ionic strength; thus, 0 and 120 mM added KCl in experiment correspond to the 8 and 128 mM simulation conditions. Experimental $\langle E \rangle$ values were measured directly for ProT$\alpha$–H1 at 8 mM ($0.45 \pm 0.03$),[16] ProT$\alpha$–H1 at 128 mM ($0.45 \pm 0.03$),[15] and ProT$\alpha$–protamine at 128 mM ($0.54 \pm 0.03$).[16] For ProT$\alpha$–protamine at 8 mM, $\langle E \rangle$ was extrapolated using an unweighted quadratic least-squares fit to all six measured salt concentrations,[16] giving $\langle E \rangle = 0.56$. The extrapolation uncertainty (0.014) was combined in quadrature with the experimental uncertainty of 0.03, giving 0.04. For comparison, the closest measured value is $\langle E \rangle = 0.56 \pm 0.03$ at 25 mM added KCl (approximately 33 mM total ionic strength). Simulation uncertainties in $\langle E \rangle$ are standard deviations across the 96 chain-specific mean transfer efficiencies and therefore report chain-to-chain heterogeneity rather than uncertainty in the condensate mean. If the 96 chain-specific mean transfer efficiencies were treated as independent samples, the corresponding standard error of the mean would be $\mathrm{SD}/\sqrt{96} \simeq 0.10\,\mathrm{SD}$. Experimental $\tau_r$ values were measured by nsFCS. For ProT$\alpha$–protamine at 8 mM, the closest experimental $\tau_r$ was measured at 25 mM added KCl (approximately 33 mM total ionic strength), because lower-salt measurements were not feasible.[16] Simulations used Amber ff99SBws ("Original") and its Lennard–Jones-refined variant ("LJ-refined"). Simulation uncertainties in $\tau_r$ are defined in Methods.

| Condensate | KCl (mM) | | $\langle E \rangle$ | $\tau_r$ ($\mu$s) |
|---|---|---|---|---|
| ProT$\alpha$–H1 | 128 | Experiment | $0.45 \pm 0.03$ | $0.38 \pm 0.04$ |
| | | Original | $0.46 \pm 0.18$ | $0.77 \pm 0.08$ |
| | | LJ-refined | $0.42 \pm 0.14$ | $0.42 \pm 0.06$ |
| ProT$\alpha$–H1 | 8 | Experiment | $0.45 \pm 0.03$ | $0.80 \pm 0.12$ |
| | | Original | $0.43 \pm 0.27$ | $0.74 \pm 0.14$ |
| | | LJ-refined | $0.43 \pm 0.16$ | $0.79 \pm 0.05$ |
| ProT$\alpha$–protamine | 128 | Experiment | $0.54 \pm 0.03$ | $2.5 \pm 0.9$ |
| | | Original | $0.50 \pm 0.33$ | $6.4 \pm 2.0$ |
| | | LJ-refined | $0.50 \pm 0.25$ | $2.8 \pm 0.3$ |
| ProT$\alpha$–protamine | 8 | Experiment | $0.56 \pm 0.04$ | $5.4 \pm 2.0$[a] |
| | | Original | $0.47 \pm 0.33$ | $18.1 \pm 2.8$ |
| | | LJ-refined | $0.54 \pm 0.25$ | $5.4 \pm 0.8$ |

[a]Experimental $\tau_r$ measured at 25 mM added KCl (approximately 33 mM total ionic strength in TEK buffer) and used for comparison with the ProT$\alpha$–protamine simulation at 8 mM KCl.

### References


(1) Bonner, O. D. Osmotic and activity coefficients of sodium and potassium glutamate at 298.15 K. *J. Chem. Eng. Data* **1981**, *26*, 147–148.

(2) Bonner, O. D. Osmotic and activity coefficients of some amino acids and their hydrochloride salts at 298.15 K. *J. Chem. Eng. Data* **1982**, *27*, 422–423.

(3) Robinson, R. A.; Stokes, R. H. *Electrolyte solutions*; Courier Corporation, 2002.

(4) Miller, M. S.; Lay, W. K.; Elcock, A. H. Osmotic pressure simulations of amino acids and peptides highlight potential routes to protein force field parameterization. *J. Phys. Chem. B* **2016**, *120*, 8217–8229.

(5) Ivanović, M. T.; Bruetzel, L. K.; Shevchuk, R.; Lipfert, J.; Hub, J. S. Quantifying the influence of the ion cloud on SAXS profiles of charged proteins. *Phys. Chem. Chem. Phys.* **2018**, *20*, 26351–26361.

(6) Lukasheva, N.; Tolmachev, D.; Martinez-Seara, H.; Karttunen, M. Changes in the local conformational states caused by simple Na+ and K+ ions in polyelectrolyte simulations: Comparison of seven force fields with and without NBFIX and ECC corrections. *Polymers* **2022**, *14*, 252.

(7) Mao, A. H.; Pappu, R. V. Crystal lattice properties fully determine short-range interaction parameters for alkali and halide ions. *J. Chem. Phys.* **2012**, *137*, 064104.

(8) Joung, I. S.; Cheatham III, T. E. Determination of alkali and halide monovalent ion parameters for use in explicitly solvated biomolecular simulations. *J. Phys. Chem. B* **2008**, *112*, 9020–9041.

(9) Reif, M. M.; Hünenberger, P. H. Computation of methodology-independent single-ion solvation properties from molecular simulations. IV. Optimized Lennard-Jones interac-

tion parameter sets for the alkali and halide ions in water. *J. Chem. Phys.* **2011**, *134*, 144104.

(10) Vanommeslaeghe, K.; MacKerell Jr, A. CHARMM additive and polarizable force fields for biophysics and computer-aided drug design. *Biochim. Biophys. Acta, Gen. Subj.* **2015**, *1850*, 861–871.

(11) Jensen, K. P.; Jorgensen, W. L. Halide, ammonium, and alkali metal ion parameters for modeling aqueous solutions. *J. Chem. Theory Comput.* **2006**, *2*, 1499–1509.

(12) Aqvist, J. Ion-water interaction potentials derived from free energy perturbation simulations. *J. Phys. Chem.* **1990**, *94*, 8021–8024.

(13) Horinek, D.; Mamatkulov, S. I.; Netz, R. R. Rational design of ion force fields based on thermodynamic solvation properties. *J. Chem. Phys.* **2009**, *130*, 124507.

(14) Martinsen, J. H.; Saar, D.; Fernandes, C. B.; Schuler, B.; Bugge, K.; Kragelund, B. B. Structure, dynamics, and stability of the globular domain of human linker histone H1.0 and the role of positive charges. *Protein Sci.* **2022**, *31*, 918–932.

(15) Galvanetto, N.; Ivanović, M. T.; Chowdhury, A.; Sottini, A.; Nüesch, M. F.; Nettels, D.; Best, R. B.; Schuler, B. Extreme dynamics in a biomolecular condensate. *Nature* **2023**, *619*, 876–883.

(16) Galvanetto, N.; Ivanović, M. T.; Del Grosso, S. A.; Chowdhury, A.; Sottini, A.; Nettels, D.; Best, R. B.; Schuler, B. Material properties of biomolecular condensates emerge from nanoscale dynamics. *Proc. Natl. Acad. Sci. U.S.A.* **2025**, *122*, e2424135122.
